# Novel antibodies detect nucleocytoplasmic O-fucose in protist pathogens, cellular slime molds, and plants

**DOI:** 10.1101/2024.10.15.618526

**Authors:** Megna Tiwari, Elisabet Gas-Pascual, Manish Goyal, Marla Popov, Kenjiroo Matsumoto, Marianne Grafe, Ralph Graf, Robert S. Haltiwanger, Neil Olszewski, Ron Orlando, John Samuelson, Christopher M. West

## Abstract

Cellular adaptations to change often involve post-translational modifications of nuclear and cytoplasmic proteins. An example found in protists and plants is the modification of serine and threonine residues of dozens to hundreds of nucleocytoplasmic proteins with a single fucose (O-Fuc). A nucleocytoplasmic O-fucosyltransferase (OFT) occurs in the pathogen *Toxoplasma gondii*, the social amoeba *Dictyostelium*, and higher plants, where it is called Spy because mutants have a spindly appearance. O-fucosylation, which is required for optimal proliferation of *Toxoplasma* and *Dictyostelium*, is paralogous to the O-GlcNAcylation of nucleocytoplasmic proteins of plants and animals that is involved in stress and nutritional responses. O-Fuc was first discovered in *Toxoplasma* using *Aleuria aurantia* lectin, but its broad specificity for terminal fucose residues on N- and O-linked glycans in the secretory pathway limits its use. Here we present affinity purified rabbit antisera that are selective for the detection and enrichment of proteins bearing fucose-O-Ser or fucose-O-Thr. These antibodies detect numerous nucleocytoplasmic proteins in *Toxoplasma, Dictyostelium*, and *Arabidopsis*, as well as O-Fuc occurring on secretory proteins of *Dictyostelium* and mammalian cells, although the latter are frequently blocked by further glycosylation. The antibodies label *Toxoplasma*, *Acanthamoeba*, and *Dictyostelium* in a pattern reminiscent of O-GlcNAc in animal cells including nuclear pores. The O-fucome of *Dictyostelium* is partially conserved with that of *Toxoplasma* and is highly induced during starvation-induced development. These antisera demonstrate the unique antigenicity of O-Fuc, document conservation of the O-fucome among unrelated protists, and will enable the study of the O-fucomes of other organisms possessing OFT-like genes.

**IMPORTANCE:** O-fucose, a form of mono-glycosylation on serine and threonine residues of nuclear and cytoplasmic proteins of some parasites, other unicellular eukaryotes, and plants, is understudied because it is difficult to detect owing to its neutral charge and lability during mass spectrometry. Yet the O-fucosyltransferase enzyme (OFT) is required for optimal growth of the agent for toxoplasmosis, *Toxoplasma gondii*, and an unrelated protist, the social amoeba *Dictyostelium discoideum*. Furthermore, O-fucosylation is closely related to the analogous process of O-GlcNAcylation of thousands of proteins of animal cells, where it plays a central role in stress and nutritional responses. O-Fuc is currently best detected using *Aleuria aurantia* lectin (AAL), but in most organisms AAL also recognizes a multitude of proteins in the secretory pathway that are modified with fucose in different ways. By establishing the potential to induce highly specific rabbit antisera that discriminate O-Fuc from all other forms of protein fucosylation, this study expands knowledge about the protist O-fucome and opens a gateway to explore the potential occurrence and roles of this intriguing posttranslational modification in bacteria and other protist pathogens such as *Acanthamoeba castellanii*.

## INTRODUCTION

Successful infection of humans by an infectious agent requires that the pathogen adapts to drastic environmental changes as it propagates and travels through its host. Post-translational modifications of proteins play crucial roles is regulating protein activity and stability. One type of post-translational modification of interest is glycosylation, in which glycosyltransferase pathways generate covalently linked oligosaccharides (glycans) to side chains of hydroxyamino acids or Asn. Most glycosylation occurs in the secretory pathway in the form of complex oligosaccharides, as summarized for mammalian cells (1,2) and evolutionary relatives in the protist kingdom (3). Within the cytoplasm and nucleus however, glycosylation typically occurs as a single monosaccharide glycosidically linked to hydroxyamino acids, though more complex examples may also occur (4).

O-fucose (O-Fuc) is a monosaccharide modification of the side chains of Ser and Thr found exclusively on nuclear and cytoplasmic proteins of the pathogen *Toxoplasma gondii* (5), plants including *Arabidopsis thaliana* (6), and the model organism *Dictyostelium discoideum* (7). O-Fuc is similar to O-GlcNAc, which modifies nucleocytoplasmic proteins in animals and plants, differing only in sugar type (8). In fact, the O-fucosyltransferase (OFT) of plants was initially misidentified as an O-GlcNAc transferase (OGT) on account of the high degree of similarity of the protein sequences (9). The relatively recent identification of the *Arabidopsis* paralog, known as Spindly or Spy, as an OFT (10) allowed attribution of previous Spy mutant phenotypes, involving hormonal control of gene expression, to O-fucosylation. The parallel discovery that homologs in *Toxoplasma* (11) and *Dictyostelium* (7) are also OFTs led to studies showing that their OFTs were required for optimal growth but not essential. Further studies showed that OFT is involved in normal dissemination of parasites to the brains of infected mice and maintenance of normal levels of several proteins (12).

O-Fuc was first discovered (5) using *Aleuria aurantia* lectin (AAL), which binds to terminal fucose wherever it occurs. Its detection in *Toxoplasma* was facilitated by the paucity of fucose on secreted proteins which, where it does occur, is masked by the addition of capping sugars (13,14). Its detection in *Dictyostelium*, which expresses over a dozen fucosyltransferases in its secretory pathway (15,16) and applies fucose as a terminal modification as is typical in most systems, required cell fractionation to isolate a cytosolic fraction from a GDP-Fuc transporter mutant to minimize contamination from the secretory pathway (7). The use of AAL in fluorescence imaging indicates a high concentration of O-Fuc in assemblies associated with the nuclear envelope of *Toxoplasma* and possibly *Dictyostelium* (5,7). Its use for pulldowns analyzed by mass spectrometry revealed dozens of O-Fuc proteins in *Toxoplasma* that were validated by detection of fucopeptides (5). More recently and in conjunction with cell fractionation, dozens of O-Fuc proteins are indicated in *Dictyostelium* (7) and hundreds in *Arabidopsis* (17,18). O-Fuc is highly enriched in intrinsically disordered regions, and favors stretches rich in Ser and Thr (5,17). The complexity in experimental design for the study of the O-fucome in fucose-rich species poses a bottleneck for the use of known lectins.

Interest in O-Fuc is motivated by its homology with O-GlcNAc, both in terms of being a monosaccharide modification of disordered regions and the homology of its glycosyltransferases, which includes around a dozen TPR domains linked to a C-terminal CAZy GT41 GT domain (7,19). Formally discovered in 1984, it has taken several decades for appropriate research tools and reagents to be widely accessible (20). O-GlcNAc is essential in animals and is now known to be reversibly applied to thousands of nucleocytoplasmic proteins (21) and to be globally involved in cellular responses to varied stresses including nutrient deprivation (8,22). O-GlcNAc acts at multiple levels of cellular regulation including transcription, nuclear transport, protein translation, protein degradation, and protein allostery. The potential for O-Fuc to have similarly extensive roles in unicellular eukaryotes is unclear but, as for O-GlcNAc, requires new reagents to facilitate investigations.

Several chemical and enzymatic methods have been developed to characterize the O-GlcNAcome (20,23) but most are not directly applicable to O-Fuc. Characterization of the O-GlcNAcome was facilitated by enrichment with plant lectins such as wheat germ agglutinin (24,20), and greatly improved by the development of monoclonal rabbit antibodies to synthetic peptide libraries containing O-GlcNAc (25). Borrowing from this example, we present here highly specific anti-O-Fuc antisera against combinatorial fucopeptide libraries containing either Fuc-O-Ser (FOS) or Fuc-O-Thr (FOT) based on flanking amino acids found in *Toxoplasma.* We document their superior ability to image O-Fuc proteins in cells at high resolution and characterize the O-fucome in fucose-rich organisms like *Dictyostelium*, the pathogen *Acanthamoeba*, and the higher plant *Arabidopsis*, and reveal aspects of the conservation and dynamics of O-Fuc.

## RESULTS

### Generation of O-Fuc-Ser (FOS) and O-Fuc-Thr (FOT) polyclonal antibodies

To develop antibodies that specifically recognized Fuc-O-Serine (FOS) in the context of a polypeptide, rabbits were immunized with a library of dodecapeptide conjugates with FOS in the seventh position (Fig. 1B). The first amino acid was Cys for conjugation, and each other position consisted of any of 10 amino acids that represent the neighboring amino acids from 33 fucopeptides sequenced from *T. gondii* (5). The library was incorporated into a proprietary conjugate optimized to stimulate the immune response. From each of 4 rabbits, successive bleeds after boosting were monitored using an ELISA method targeting the fucopeptide peptide library or a corresponding peptide library conjugated to BSA. As described in Fig. 1A, the antisera were negatively absorbed against immobilized naked peptides and affinity purified against immobilized FOS peptides, before being used in assays to describe the O-fucome in various organisms. A representative purified antiserum yielded a titer against FOS peptides up to 1:100,000 and minimal reactivity toward Ser-peptides (Fig. 1C). A parallel strategy yielded affinity purified antisera selective for FOT-peptides (Fig. 1D).

**Fig. 1.**
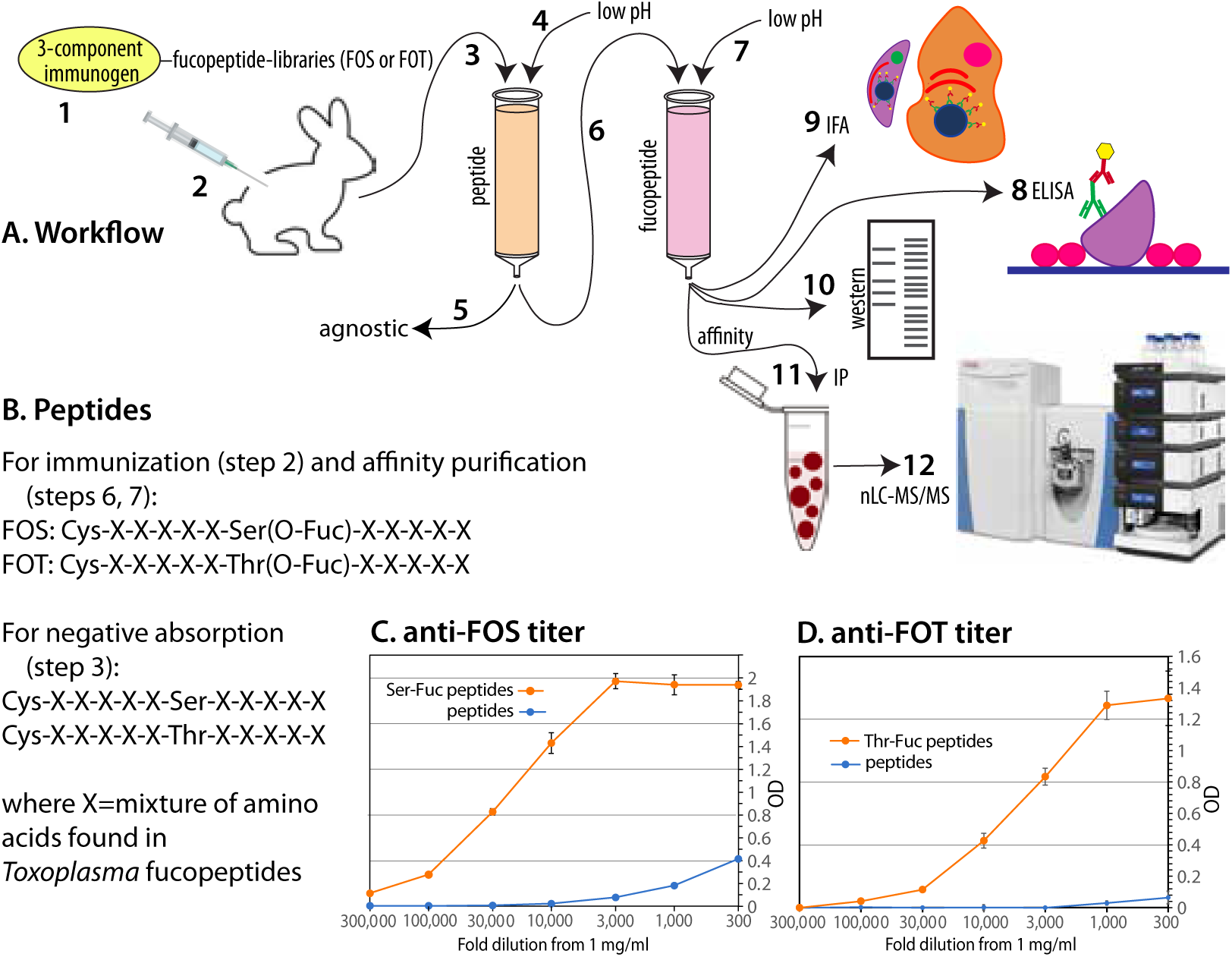
Anti-fucopeptide antisera. (A) Workflow of antibody generation and application to O-fucome studies. The fucopeptide library immunogens (1) were injected into rabbits (2). After boosting, serum was collected and passed over a peptide library column (3), which was eluted with low pH buffer (4) to produce an agnostic reference antiserum (5). The flow-through from the peptide column was applied to a fucopeptide library column (6), which was eluted with low pH (7). The resulting affinity purified antiserum was assayed by ELISA(8), and used for immunofluorescence analysis of cells (9), western blotting of cell extracts (10), and for affinity capture of O-Fuc proteins (11) for proteomic analysis using nLC-MS/MS (12). (B) Description of the fucopeptide libraries, containing FOS or FOT, and peptide libraries containing Ser or Thr, that were incorporated via their N-terminal Cys residues into a tripartite immunogen (1), to columns for negative or positive affinity purification (3, 6), to BSA for ELISA assays (8), or to beads for affinity capture (11). (C) A dilution series of a representative bleed of affinity purified anti-FOS antisera was assayed for binding to the FOS library or the unmodified Ser-peptide library, using an ELISA method based on alkaline phosphatase conjugated goat-anti-rabbit IgG. (D) Titration of a representative affinity-purified bleed a rabbit immunized with the FOT peptide library. Error bars represent S.D. of 2 technical replicates.

### Anti-O-FOS/T antibody specificity based on immunofluorescence microscopy of *Toxoplasma*

The specificities of anti-FOS peptide and anti-FOT peptide (referred to anti-FOS and anti-FOT, or as anti-FOS/T when generalizing) were examined using indirect immunofluorescence analysis (IFA) of fixed wild-type intracellular tachyzoites of *Toxoplasma* grown in human foreskin fibroblasts (HFFs) (Fig. 2). Pilot studies to optimize the method favored methanol over paraformaldehyde-fixation, 0.3-1 µg/ml IgG concentration, and blocking with animal serum (see Methods). To visualize parasites within host parasitophorous vacuoles, samples were labeled with antibodies to the cell surface protein SAG1. Based on super-resolution structured illumination microscopy (SR-SIM), anti-FOS staining was mostly associated with punctae in parasite nuclei, with greater density near the nuclear periphery (Fig. 2A), similar to the staining pattern observed with AAL (Fig. 2B). A lower level of diffuse nucleocytoplasmic staining cannot be excluded owing the normalization method used for SR-SIM. Secondary Ab and streptavidin controls were negative (not shown). Preincubation with the hapten inhibitor α-methyl fucose (αMeFuc) blocked labeling indicating specificity of anti-FOS toward fucose. TgSPYΔ parasites were not labeled (Fig. 2A) confirming dependence on O-Fuc conjugated to nucleocytoplasmic proteins. Furthermore, HFF host cells were completely devoid of staining with anti-FOS, in contrast to robust labeling of host cells with AAL. This is consistent with absence of SPY and known nucleocytoplasmic O-Fuc in mammalian cells and indicates that anti-FOS is selective for Fuc linked to amino acids rather than as a terminal modification of N- and O-glycans that is abundant on mammalian but not parasite glycans. Results with anti-FOT were essentially indistinguishable from those with anti-FOS (Fig. S1A). As expected, probing with agnostic pAbs (Fig. 1A) yielded no distinct pattern of labeling (Fig. S1B-C).

**Fig. 2.**
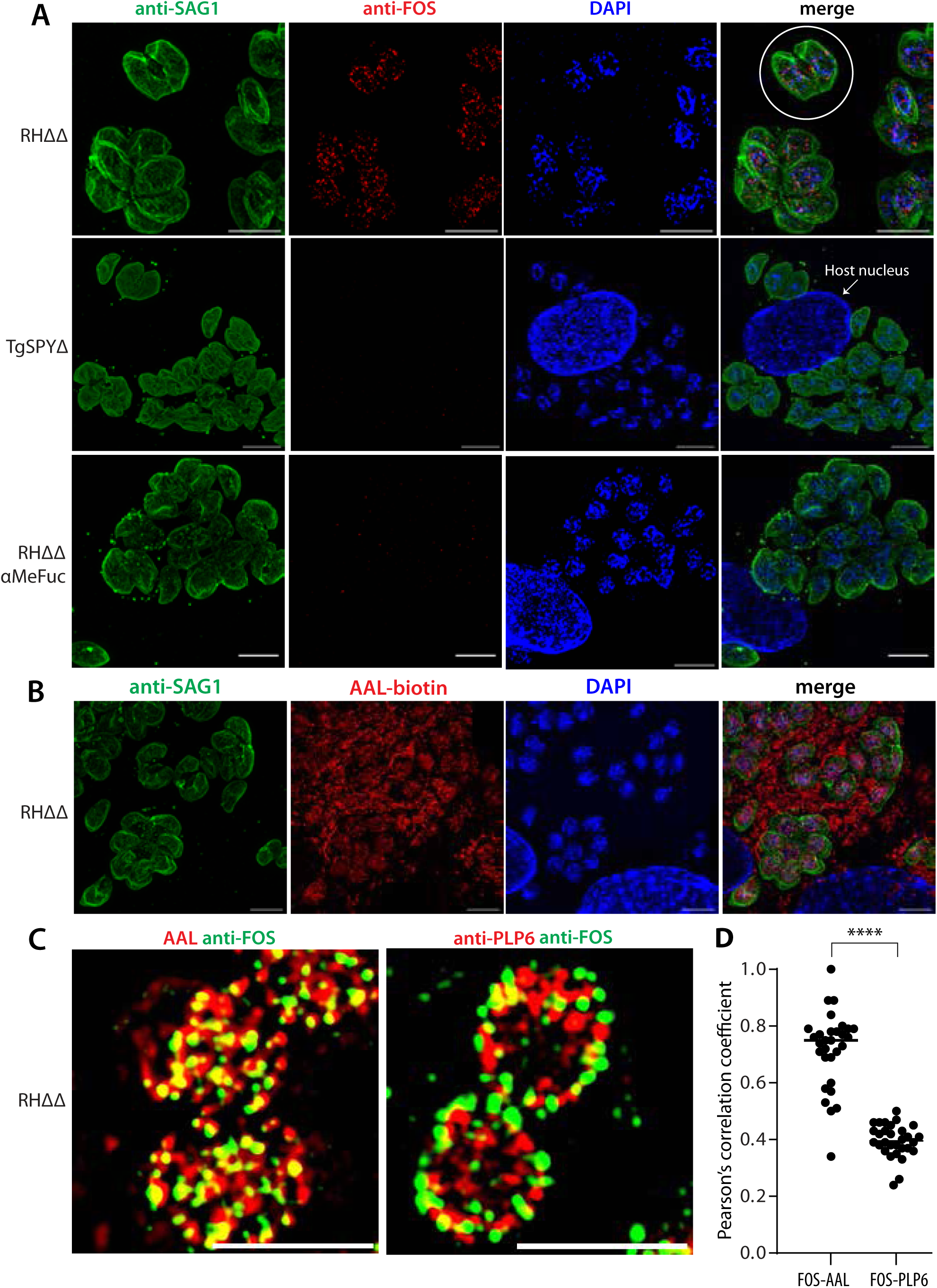
Immunofluorescence analysis of anti-FOS specificity using *Toxoplasma* infected HFFs. Monolayers of human foreskin fibroblasts were infected with parasites, fixed, and imaged using Super-Resolution Structured Illumination Microscopy (SR-SIM). (A) HFFs containing wild-type RHΔΔ or TgSPYΔ parasites were probed with affinity purified rabbit anti-FOS (1 µg/ml) and mouse anti-SAG1 followed by Alexa Fluor-488 goat anti-rabbit IgG and Alexa Fluor-594 goat anti-mouse IgG to localize O-Fuc and outline the parasites, respectively. DAPI (blue) was used to visualize parasite and host cell nuclei (an example is labeled). An instance of 2 parasites occupying a parasitophorous vacuole within an HFF is outlined with a white circle. In the lower row, RHΔΔ samples were probed with anti-FOS in the presence of 0.2 M αMeFuc. Maximum intensity projections are shown. (B) Same as panel A, except that 1 µg/ml AAL-biotin and Alexa Fluor-594 streptavidin were used in place of anti-FOS. Scale bars: 5 µm. See Fig. S1 for corresponding probing with anti-FOT. (C) Same as above, except that samples were probed with anti-FOS and either AAL-biotin or anti-PLP6 (1:5000), a marker of nuclear epichromatin. Representative single z-slices of the nuclear regions are shown. See Fig. S2 for additional images with anti-FOS/T. (D) Pearson’s correlation coefficients for colocalization probes in panel C. Each point represents a single cell (n>30), and means are represented with a horizontal bar. The difference was significant using an unpaired, two-tailed t-test (****p< 0.0001). Scale bars: 5 µm.

In co-labeling experiments, anti-FOS localized in close proximity to, tending in some cases to encircle, an epichromatin epitope that is enriched under the nuclear envelope (26,27) and recognized by mAb PLP6 (Fig. 2C). Co-labeling of anti-FOS with AAL-biotin showed that, as expected, both were predominantly nuclear with considerable overlap (yellow) of punctae (Fig. 3C) at the maximal lateral (x-y) resolution of ∼100 nm and ∼ 200 nm axial (z) resolution of the microscope system. Comparison of the Pearson’s correlation coefficients (PCC) indicated a significantly higher overlap of FOS with AAL (0.72) than with PLP6 (0.40) (Fig. 2D). Similar findings were observed in a comparison of anti-FOT and AAL (Fig. S2). The imperfect overlap of anti-FOS and AAL might be due to segregation of FOS and FOT targets or, alternatively, differences in accessibility to indirect probing with anti-FOS/T and AAL or avidity effects.

**Fig. 3.**
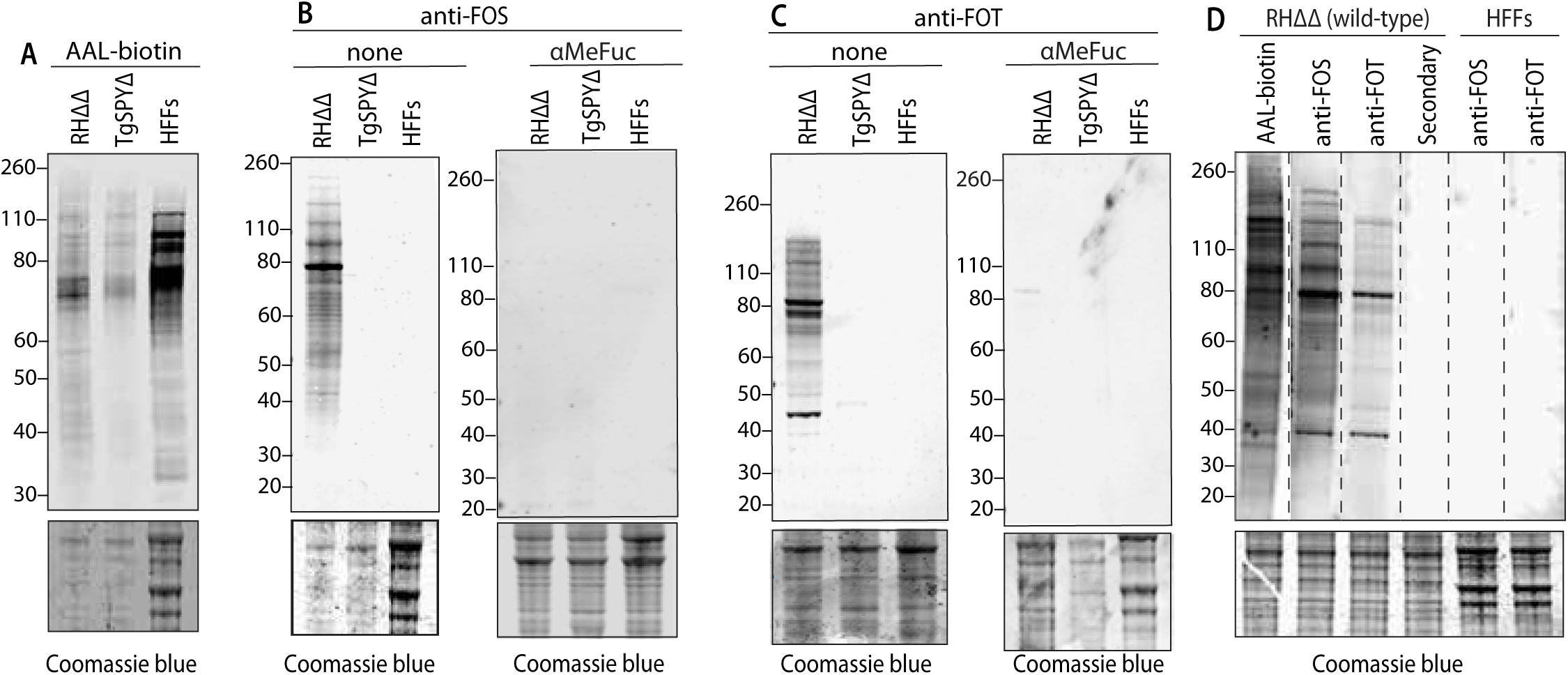
Western blot analysis of anti-FOS and anti-FOT specificity in *Toxoplasma*. Parasites were harvested from hTERT fibroblasts as a mixture of intracellular and freshly spontaneously lysed out cells. In parallel with hTERT fibroblasts, samples were dissolved in SDS and analyzed by SDS-PAGE/western blotting. Post-blot gels were stained with Coomassie blue as loading controls. (A) Samples from RHΔΔ (wild-type) and SPYΔ strains, and host cells, were probed using 1 µg/ml AAL-biotin and Alexa Fluor-680 streptavidin. (B) The same samples were probed with 1 µg/ml anti-FOS and Alexa Fluor-680 goat-anti rabbit IgG, either with or without 0.2 M αMeFuc. (C) Same as panel B using 0.3 µg/ml anti-FOT. (D) Strips from a single western blot were probed with AAL-biotin, anti-FOS, anti-FOT or no primary probe (secondary) as indicated. Dashed lines represent boundaries between strips.

### Anti-O-FOS/T specificity based on western blot analysis of *Toxoplasma* proteins

Whole cell extracts of tachyzoites harvested from HFF monolayers were western blotted to survey the range of potential O-Fuc proteins. Labeling with anti-FOS and anti-FOT was optimized at 1 µg/ml and 0.3 µg/ml, respectively. Both anti-FOS (Fig. 3B) and anti-FOT (Fig. 3C) labeled numerous proteins over the *M_r_* range of 40,000 to >260,000. Negligible bands above background were detected in the TgSPYΔ strain, nor in HFF host cells, or secondary Ab controls (Fig. 3D). This is significant because AAL, with its broader anti-Fuc specificity, labeled residual proteins in TgSPYΔ (Fig. 3A), which likely represented contamination from host cells or culture media serum, which are rich in other classes of fucoconjugates. A side-by-side comparison of the western blot profiles show similarities and differences among AAL, anti-FOS and anti-FOT (Fig. 3D). Though the complexity of the patterns complicate interpretation, dissimilarities suggest differential expression of FOS and FOT on different proteins. Differences with the AAL profile might be explained by differences in avidity and accessibility in the western blot setting, but the possibility that the Abs are not fully pan-specific cannot be excluded. Overall, the western blot studies confirm the specificity indicated by the immunofluorescence studies and draw attention to the large number of proteins that are modified with O-Fuc.

### Specificity of anti-FOS/T based on immunofluorescence analysis of *Dictyostelium*

*Dictyostelium discoideum* is a social soil amoeba belonging to the amorphoeae group that is evolutionarily closely affiliated with metazoa and far removed from the apicomplexan *Toxoplasma gondii*. This dissimilarity and its high level of fucosylation in the secretory pathway recognized by AAL led us to choose it for comparison (7). Immunofluorescence analysis of growth stage (vegetative) amoebae showed an anti-FOT labeling pattern similar to *Toxoplasma*, with signal concentrated in nuclear punctae (Fig. 4B). As above, the SR-SIM method tends to emphasize focal concentrations, so an additional more diffuse localization cannot be excluded. Comparison with AAL emphasizes the futility of using this lectin to differentiate Spy-dependent O-Fuc from other forms of fucosylation occurring in the secretory pathway and cell surface (Figs. 4A,S3A). Fig. 4B shows that anti-FOS labeling is absent in *spy*-KO amoebae outlined with anti-actin to profile the cells, confirming dependence on O-Fuc. Furthermore, labeling is not affected in *gft*-KO (*gft*^−^) cells, which are unable to fucosylate secretory and plasma membrane proteins and lipids on account of absence of the GDP-Fuc transporter (GFT) required to provide GDP-Fuc to the luminal fucosyltransferases (Fig. S3B). Co-localization of anti-FOS with anti-PLP6 reveals that the peripheral nuclear punctae of FOS and FOT reside beyond the domain of the epichromatin marker (Figs. 4C,S3C), which is more apparent in *Dictyostelium* due to its larger size relative to *Toxoplasma*. The possibility that O-Fuc punctae are associated with nuclear pores was addressed by co-localizing with DdNup62 (Nucleoporin 62, DDB_G0274587), an FG-repeat protein that resides in the central pore, and DdNup210 (Nucleoporin 210, DDB_G0288545), a transmembrane protein circumscribing the pore ring (31). Both were found in the *Dictyostelium* O-fucome (see below). For comparison, we examined co-localization with an inner nuclear membrane marker, Src1 (36), which is associated with nucleolar attachments. These proteins were previously tagged at their C-termini with mNeonGreen using a knock-in approach (31). Nup62-mNeon and Nup210-mNeon exhibited puncta-like distributions around the nucleus, as expected for nuclear pores, that were partially overlapped by anti-FOS (Fig. 4D) and anti-FOT (Fig. S3D) labeling. In contrast, Src1-mNeon localized asymmetrically as expected for co-association with nucleoli, with considerably less apparent overlap with anti-FOS or anti-FOT. Based on Pearson’s correlation coefficients (PCC) to quantify co-localization, the associations of anti-FOS and anti-FOT with Nup62-mNeon and Nup210-mNeon were confirmed to be higher than with Src1-mNeon (Fig. 4E). Interestingly, anti-FOT exhibited a significantly higher overlap with Nup62-mNeon (0.56) than did anti-FOS (0.48), which correlated with pulldown data (see below) and the presence of a poly-Thr rich tail toward the C-terminus of the protein. Thus, anti-FOS/T verified the evolutionary conservation of nuclear pore enrichment of O-Fuc proteins.

**Fig. 4.**
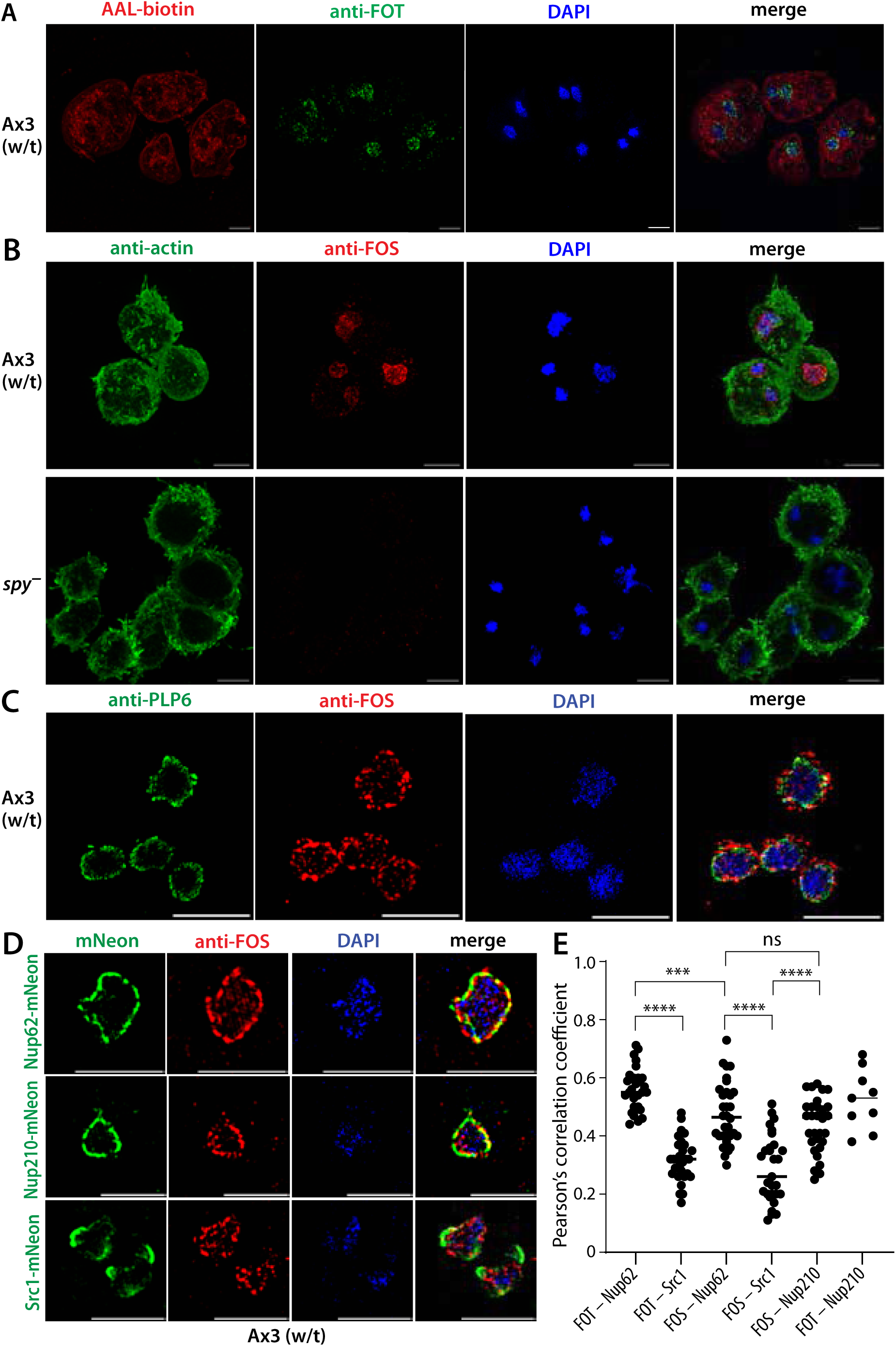
Immunofluorescence localization of anti-FOT and anti-FOS in *Dictyostelium* amoebae. Growth stage amoebae were allowed to adhere to coverslips, fixed in cold methanol, blocked in BSA, probed with the indicated reagents, and imaged with SR-SIM as in Fig. 2. (A) Wild-type (w/t, strain Ax3) amoebae were probed with anti-FOT and AAL-biotin, followed by Alexa Fluor-488 goat anti-rabbit IgG and Alexa Fluor-594 streptavidin. DAPI (blue) was used to visualize nuclei. Multinuclearity results from inefficient cytokinesis when grown in shaking culture. Maximum projection images are shown. (B) As in panel A, except that w/t and Dd*spy*-KO (*spy*^−^) amoebae were probed with anti-FOS and murine anti-actin followed by Alexa Fluor-488 goat anti-rabbit IgG and Alexa Fluor-594 goat anti-mouse IgG. See Fig. S3 for corresponding probing with anti-FOT. Scale bars: 5 µm. (C) As in panel A, except that amoebae were probed with anti-FOS and anti-PLP6 (1:5000), followed by Alexa Fluor-488 goat anti-rabbit IgG and Alexa Fluor-594 goat anti-mouse IgG. (D) Comparison of anti-FOS and anti-FOT with nuclear envelope proteins. Growth stage amoebae whose Nup62, Nup210, or Src1 genomic loci were C-terminally tagged with mNeon were probed with anti-FOS followed by Alexa Fluor-594 goat anti-rabbit IgG, and co-imaged with intrinsic mNeon fluorescence and DAPI. Single slice images are shown. See Fig. S3D for data for anti-FOT. (E) Pearson’s correlation coefficients for colocalization of anti-FOS and anti-FOT with Nup62-mNeon, Nup210-mNeon, and Src1-mNeon, calculated from super-resolution images. Each point represents a single cell, and means are indicated with a horizontal bar. Analyses were performed with ≥30 cells (except FOT – Nup210, which was not statistically evaluated). Significance was evaluated using an unpaired, two-tailed t-test (****p<0.0001, ***p<0.001, **p<0.01, *p<0.05, ns=not significant). Scale bars: 5 µm.

### Specificity of anti-FOS/T based on western blot analysis of developing *Dictyostelium*

Western blot analysis of growth stage amoebae revealed multiple proteins reactive with anti-FOS and anti-FOT, that were not observed in *spy*-KO cells (Figs. 5B,C). As expected, they were also detected in *gft*-KO cells, confirming their presence in the nucleocytoplasmic compartment. Notably, the profile was distinctive between anti-FOS and anti-FOT, suggesting differential modifications as suggested for Nup62 above. Similar to amoebae staining (Fig. 4A), AAL labeling was dominated by fucosylation in the secretory pathway as confirmed by reduced labeling in *gft*-KO cells; this reinforces the utility of anti-FOS/T to selectively detect O-Fuc (Fig. 5A). To begin to explore the possibility that O-fucosylation is developmentally regulated, samples were collected over the time course of starvation induced aggregation and fruiting body formation at 24 h. Probing the wild-type strain Ax3 with anti-FOS revealed a rather dramatic change in the *M*_r_ profile attended by a substantial increase in labeling intensity (Fig. 5D), which might be explained by increased site occupancy, a change in which proteins are O-fucosylated, and/or appearance of new O-Fuc proteins. As expected, a similar progression was observed in *gft*-KO cells. Interestingly, labeling of discrete bands were noted at later developmental times in the *spy*-KO strain but not apparent in *gft*-KO cells. This is potentially due to O-Fuc previously described on spore coat proteins (37,38), and further supported by detection of O-Fuc transferase activity in extracts (39). Interestingly, an altered profile of more intensely labeled bands appeared during development of a Spy^oe^ strain (Fig. 5D), where Spy is expressed under control of a semi-constitutive promoter (7). These changes suggest that O-fucosylation might be regulated with perhaps a hierarchical preference of substrates at different enzyme activity levels. A related though not as pronounced trend was exhibited by a more limited range of proteins using anti-FOT (Fig. 5E), at least within the sensitivity of the method. Thus applying anti-FOS/T to whole *Dictyostelium* extracts was particularly useful for mapping the extent and dynamic nature of the O-fucome during development in the context of all other fucosylation.

**Fig. 5.**
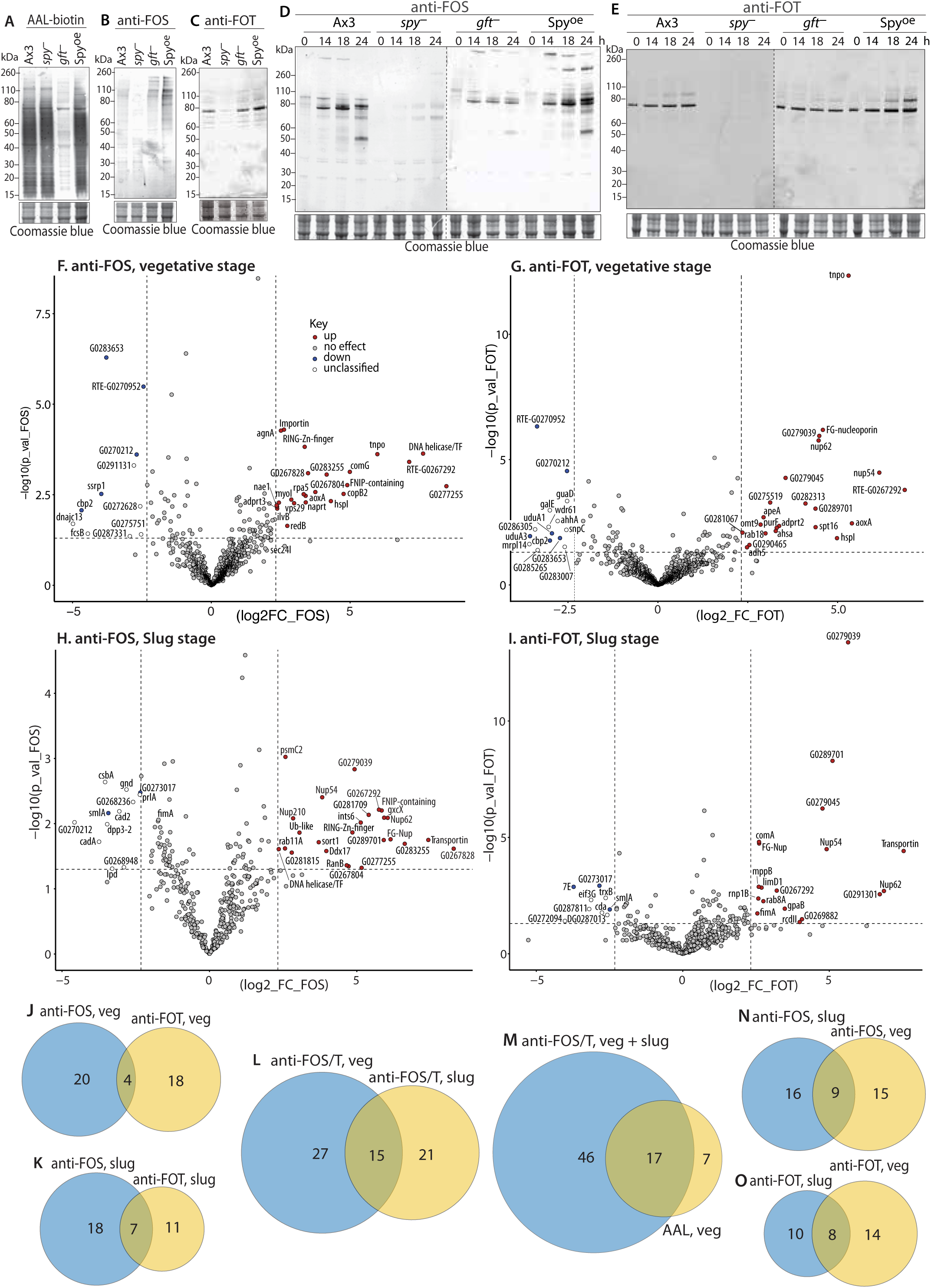
Developmental regulation of the *Dictyostelium* O-fucome. (A-C) The indicated strains of growth stage (vegetative) amoebae were subjected to SDS-PAGE, western blotting, and immunoprobing as in Fig. 3. The blots were probed with anti-FOS (A), anti-FOT (B), or AAL-biotin (C) with secondary Abs or streptavidin. Excerpts from the Coomassie blue stained blotted gels are shown as loading controls. (D, E). Growth stage amoebae (0 h) were deposited on non-nutrient agar and harvested after the indicated number of h for western blotting with anti-FOS (D) or anti-FOT (E). Multicellular slugs are formed by 14 h, Culmination begins around 18 h, and fruiting bodies are formed by 24 h. Molecular weight marker positions are shown in kDa. (F-I) Proteomics analysis of the O-fucome. Proteins were captured from CHAPS-solubilized whole cell extracts from wild-type (strain Ax3) or Dd*spy*-KO cells with anti-FOS or anti-FOT cross-linked to Protein A/G magnetic beads. Proteins eluted with αMeFuc in 8 M urea were reduced and alkylated, converted to peptides using endo Lys-C and trypsin, and analyzed by nLC using an Orbitrap mass analyzer. Proteins were identified based on detection of ≥2 peptides (1% FDR) at <5% protein FDR. The Volcano plots report on the fold enrichment in wild-type vs. *spy*-KO cells (abscissa) vs. confidence of fold-change for 3 biological replicates each with 3 technical replicates for a total of 9 data pairs for each protein. Data from anti-FOS and anti-FOT pulldowns of vegetative stage extracts are shown in panels F and G, and similarly for slug stage extracts in panels H and I. The ‘DDB_’ prefix is removed from the labels in Table 2, which lists wild-type enriched proteins (upper red quadrant, in red) along with which fractions they were found. See Table S1 for an analysis of proteins over-enriched in *spy*-KO extracts (upper left quadrant). Proteins for whose over-enrichment was supported in complementary pulldowns are in blue, and those without support are in white (see text for explanation). (J-O) Summary of O-Fuc proteins detected. Venn diagrams show distribution of O-Fuc candidates between anti-FOS and anti-FOT pulldowns, and between vegetative (veg) and slug stage cells, as indicated. Categories are illustrated in Table 2. Panel M includes candidates from a previous AAL-pulldown (7).

### Anti-FOS/T antibodies identify many O-Fuc proteins in the *Dictyostelium* developmental life cycle

The identity of the O-fucosylated proteins was examined by coupling anti-FOS and anti-FOT to magnetic beads for use in a proteomics workflow. Optimization of the ratio of pAb-beads to detergent solubilized extracts of whole amoebae resulted in near quantitative capture of major bands as detected by western blotting. nLC/MS was employed to identify proteins based on peptides detected and quantified from spectral counting according to criteria described in Methods. Volcano plots of proteins enriched in pulldowns with anti-FOS (Fig. 5F) and anti-FOT (Fig. 5G) from wild-type compared to *spy*-KO vegetative cells indicated that a 5-fold enrichment factor identified the majority of proteins found in a previous study using AAL. In contrast, the great majority of known non-specific binders, secretory pathway residents, and other organelle proteins not expected to be accessible to OFT did not achieve this enrichment level. Unexpectedly, several proteins were found to be enriched over 5-fold in the *spy*-KO strain. Since there is no evidence for an unknown neo-epitope in *spy*-KO cells, these proteins (listed in Table S1) likely bind non-specifically and are potentially increased in *spy*-KO cells. If correct, it is expected that similar negative enrichment would occur for both anti-FOS and anti-FOT. Seven proteins satisfy this criterion in vegetative cells, and only 3 different proteins do so in slug cells (Figs. 5H,I).

Together, 42 proteins were identified in anti-FOS and anti-FOT pulldowns from growing whole cell extracts (Table 2). Of these, 24 were found with anti-FOS and 22 with anti-FOT, and only 4 were found in both (Fig. 5J). In a parallel analysis of 14-h slugs, Volcano plots yielded similar findings, with 36 proteins matching the criteria for O-Fuc candidates (Figs. 5H,I). Of these, 25 were found with anti-FOS, and 18 with anti-FOT, with 7 of these were found in both (Fig. 5K). Thus anti-FOS and anti-FOT exhibit differential selectivity. Notably, only 15 of the proteins were detected in both vegetative and slug cells (Fig. 5L), which correlates with the distinct western blot profiles (Figs. 5D,E) between these two developmental stages. All but 7 of the 24 proteins captured by AAL from nucleocytoplasmic preparations from *gft*-KO growing cells (7), but not found in a *gmd*-KO (GDP-Fuc synthesis mutant), were >5-fold enriched in wild-type vs. *spy*-KO cells in all of the anti-FOS/T pulldowns (Fig. 5M). Furthermore, 3 of those 7 were detected at a lower enrichment threshold (>2-fold). Thus together the pAbs are broadly pan-specific.

The identified proteins are associated with a broad range of functions or sequence motifs (Table 2). Of the 15 proteins found in both growing and developed (slug) cells, 7 were FG-rich nuclear pore-like proteins particularly enriched in anti-FOT pulldowns, consistent with the cellular localization studies. Other proteins were associated with nuclear transport, transcription, and the ubiquitin-proteasome system. Stage specific proteins included additional members of these classes, as well as proteins associated with DNA repair, RNA processing, translation, stress responses, signaling, cytoskeletal network, membrane trafficking, and numerous metabolic enzymes. Two proteins expected to occur within the mitochondria might have been modified before import, though the possibility of non-specific capture of a protein underexpressed in *spy*-KO cells cannot be excluded. The number of proteins detected are comparable between *Dictyostelium* and *Toxoplasma*, though a recent deep analysis of the *Arabidopsis* O-fucome suggests that hundreds of proteins are O-fucosylated (17). Although there is not enough information to map these proteins directly to the reported O-fucomes of *Toxoplasma* (5), *Arabidopsis* (17), and *Nicotiana* (18), the protein categories largely overlap.

The stepped CID methods used to identify the peptides by mass spectrometry tend to detach the Fuc precluding direct confirmation of its presence and mapping its sites. However, using MS-MS, O-Fuc was partially mapped on two peptides from an FG-repeat nucleoporin and a peptide from a putative helicase/transcription factor, from anti-FOS pulldowns (Figs. S4,5). The sequences of these 3 peptides (STSPVTTSIP*SFTSSSTSTS*LFGPTTTTTDKDSK, TDFIGINGV*STSTSSSTSTS*K, STIIP*SSSSSSSSSS*SSSSSSSSSSTQSK, potential O-Fuc site in italics) are consistent with other evidence indicating a strong preference for Ser-, Thr-, and Ser/Thr-rich tracts (5,17). Notably, the presence of one or two copies of O-Fuc was also mapped to an anti-FOT peptide (TDEECCPIIPICINP*STIAASTIATTTAST*R) from slug extracts, assigned to the spore coat protein SP70 (Fig. S6). SP70 is a secretory protein that would not be expected to be accessible to OFT, and indeed its presence in the extracts was not enriched in wild-type vs. *spy*-KO slug cells. This finding is consistent with the Western blot evidence for anti-FOS and anti-FOT reactivity at the position of SP70 in *spy*-KO slugs and fruiting bodies (Figs. 5D,E). These pAbs confirm the presence of this O-Fuc on SP70, and demonstrate their utility for detecting non-OFT-mediated O-Fuc. Thus the new pAbs allowed an expansion of the known O-fucome, achieving 70 overall candidates including most of those detected using AAL. Furthermore, they were able to substantially differentiate between proteins enriched in FOS or FOT, validate the developmental regulation of O-Fuc in *Dictyostelium* inferred from western blotting, and identify Spy-independent O-Fuc.

### Specificity of anti-FOS/T in the pathogenic amoeba *Acanthamoeba castellanii*, the higher plant *Arabidopsis thaliana* and mammalian secretory proteins

*Acanthamoeba* is a protist pathogen belonging to the amoebozoa group with *Dictyostelium*. *A. castellanii* is a ubiquitous soil-dwelling protist with the capacity to encyst in a stable dormant state (40) and reactivate to cause irreparable damage to the cornea following contamination of contact lenses (41–43). *Acanthamoeba* exhibited substantial fucosylation throughout its secretory and endolysosomal pathways and cell surface as evidenced by AAL staining of the trophozoite stage (Fig. 6A), making it impossible to verify the presence of O-Fuc that is predicted by the presence of an OFT-like sequence in its genome. In contrast, labeling with anti-FOS or anti-FOT was highly enriched in the nucleus. This was reminiscent of the pattern observed in *Dictyostelium* and confirmed the utility of anti-FOS/T for a fucosylation-rich pathogen.

**Fig. 6.**
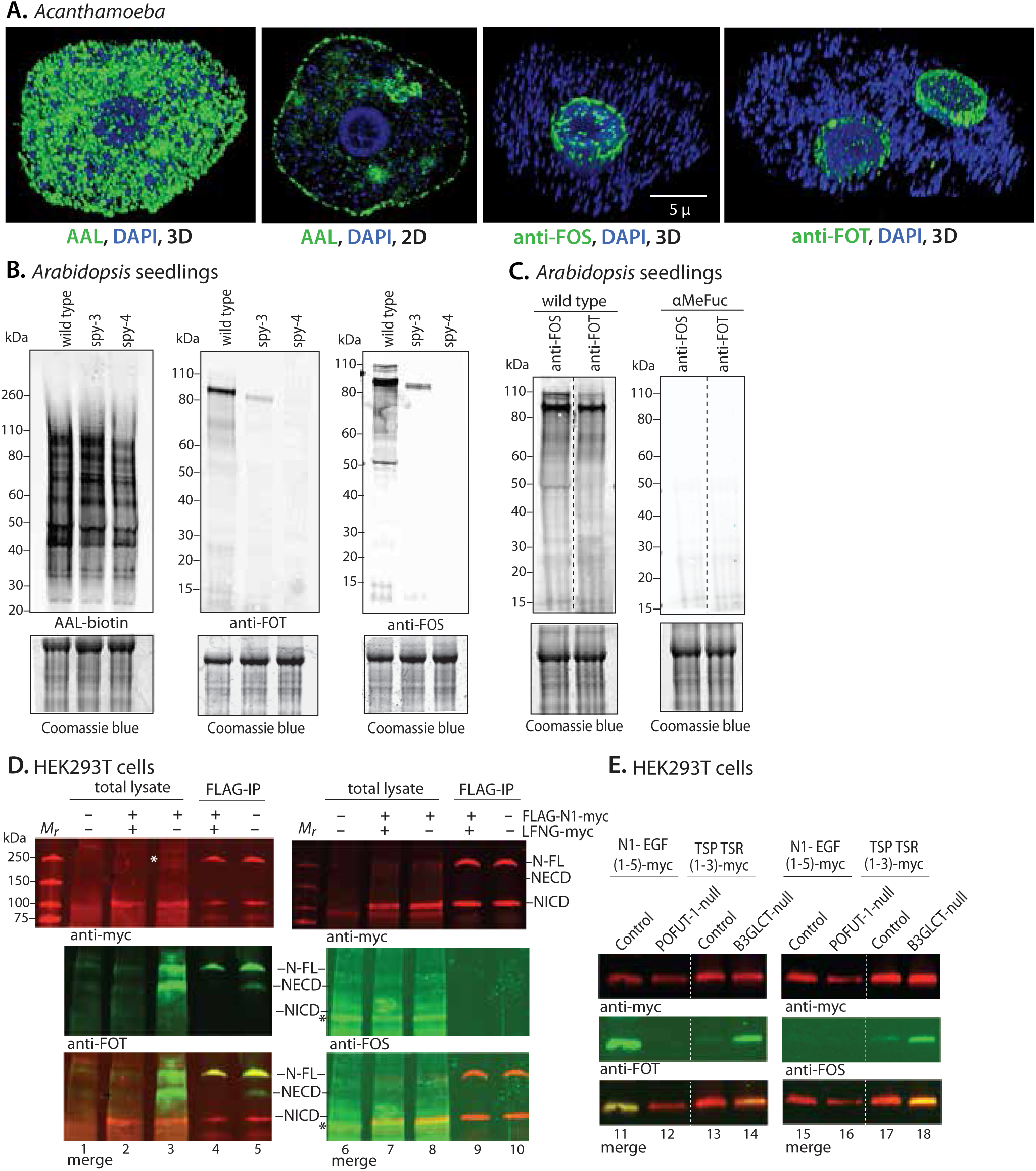
Recognition of O-Fuc proteins from *Acanthamoeba*, *Arabidopsis*, and mammalian cells. (A) Localization of AAL, anti-FOS and anti-FOT in *Acanthamoeba castellanii*, based on confocal microscopy. 2D=optical section; 3D=maximum projection). Note: DAPI labels nuclei and hundreds of mitochondria, which have an unusually large genome. (B,C) Western blot analysis of *Arabidopsis* seedlings. Protein was extracted from whole plant seedlings of wild-type (Columbia) and spy-3 and spy-4 strains, western blotted, and probed with AAL-biotin, anti-FOS, or anti-FOT, followed by streptavidin or secondary Abs as above. The post-blot gels were stained with Coomassie blue as loading controls. In panel C, w/t proteins were run on the same gel and slices were probed with anti-FOS or anti-FOT, with or without αMeFuc. Vertical dashed lines indicate slicing of western blots for probing with different Abs. Positions of molecular weight markers are shown in kDa. (D,E) anti-FOT and anti-FOS were used to detect O-Fuc on mammalian Notch-1 and Thrombospondin-1. HEK293T cells were transiently transfected to overexpress epitope-tagged human Notch1 (FLAG-N1-myc), and co-transfected with the myc-tagged LFNG capping glycosyltransferase as indicated. FLAG- and myc-tags are on the extracellular and intracellular domains of FLAG-N1-myc, respectively. Western blots of total cell extracts, or material captured by immunoprecipitation with anti-FLAG (FLAG-IP), were probed with anti-myc for FLAG-N1-myc, or with anti-FOT or anti-FOS for O-Fuc, followed by Alexa Fluor-800 or Alexa Fluor-680 fluorescent secondary Abs for the protein or O-Fuc, respectively (D). Partial processing of FLAG-N1-myc generated the myc-tagged intracellular domain (NICD) and the fucosylated extracellular domain (NECD). At high laser scanning intensity used to search for anti-FOS labeling, non-specific binding was observed (*). A truncated version of N1 consisting of 5 EGF repeats, or truncated TSP1 with 3 TSR repeats, both epitope-tagged, were transiently overexpressed in normal, POFUT1-null or B3GLCT-null HEK293T cells (E). Secreted truncated proteins were captured from culture media with anti-myc Ab and subjected to western blot analysis as in panel D.

Recently, several hundred nucleocytoplasmic targets of AtSPY were identified utilizing AAL (17,18), with many of the identified substrates being similar to OFT targets identified in *Toxoplasma* and *Dictyostelium* (5,7). Since the *Arabidopsis* studies required laborious prior cell fractionation to enrich for cytosol, we tested the application of anti-FOS/T in w/t and two homozygous hypomorphic SPY mutant strains (Table 1). Western blotting of whole seedling plant extracts with anti-FOS detected multiple bands that were a small subset of proteins that bound AAL (Fig. 6B). Inhibition by αMeFuc confirmed specificity for fucose (Fig. 6C). Reactivity was strongly reduced in the spy-3 strain (which has a missense mutation), and even further reduced in the spy-4 strain (which has a T-DNA insertion in its promoter). This was consistent with spy-4 plants being phenotypically much more severe than spy-3, with greatly reduced but detectable RNA levels by RT-PCR indicating spy-4 as not null. A similar trend was observed with the anti-FOT antibody. Side-by-side analysis of wild-type seedlings probed with anti-FOS and anti-FOT showed minor differences suggesting distinct targets for each (Fig. 6C). Collectively these results indicate that anti-FOS/T antibodies are specific for SPY-dependent modifications in *A. thaliana* and confirms that the majority of AAL-targets are unrelated to O-Fuc.

**Table 1.** Cells used in this study.

| Strain name | Parental strain | Genotype | Selection marker | Reference |
| --- | --- | --- | --- | --- |
| ***Toxoplasma gondii*** |  |  |  |  |
| RH$\Delta\Delta$ (type 1) | RH | $\Delta ku80/\Delta hxgp rt$ | None | 28 |
| RH$\Delta\Delta$SPYΔ | RH$\Delta\Delta$ | $\Delta ku80/\Delta hxgp rt/\Delta spy$ | HXGPRT | Tiwari et al., unpublished |
| ***Dictyostelium discoideum*** |  |  |  |  |
| Ax3 | NC-4 | *axeA/B/C* | None | 29 |
| *spy*-KO | Ax3 | $\Delta spy-2a$ | blasticidin-S | 7 |
| *spy*<sup>oe</sup> | Ax3 | *dscC::spy (oe-1)* | G418 | 7 |
| *gft*-KO | Ax3 | $\Delta gft-2$ | blasticidin-S | 7 |
| Ax2 | NC-4 | *axeA/B/C* | None | 30 |
| Nup62-mNeon | Ax2 |  | blasticidin-S | 31 |
| Nup210-mNeon | Ax2 |  | blasticidin-S | 31 |
| Src1-mNeon | Ax2 |  | blasticidin-S | 31 |
| ***Acanthamoeba castellanii*** |  |  |  |  |
| Neff strain | wild-type |  |  | ATCC 30010 |
| ***Arabidopsis thaliana*** |  |  |  |  |
| Columbia | wild-type |  |  |  |
| *spy*-3 | Columbia |  |  | 32 |
| *spy*-4 | Columbia |  |  | 33 |
| ***Homo sapiens*** |  |  |  |  |
| HFF |  |  |  | ATCC |
| hTERT |  |  |  | ATCC |
| HEK293T |  |  |  | ATCC |
| HEK293T-POFUT1<sup>-/-</sup> |  |  |  | 34 |
| HEK293T-B3GLUT<sup>-/-</sup> |  |  |  | 35 |

Mammalian cells lack the Spy OFT and, as shown above, HFFs were consistently negative for anti-FOS/T labeling (Figs. 2A,3). However, mammalian cells do express several protein O-fucosyltransferases (POFUTs) in the rER that also modify Ser or Thr residues (44,45). These occur within sequence motifs within pre-folded domains of secretory proteins, including epidermal growth factor (EGF) repeats, thrombospondin repeats (TSR) and elastin microfibril interface (EMI) domain repeats (45,46). These examples offered an additional opportunity to explore the pan-specificity of anti-FOS/T. Western blot analysis of total cell lysates of HEK293T cells failed to specifically detect O-Fuc above background (Fig. 6D, lane 1), which could be explained by low abundance together with partial capping by another sugar. As a first step, epitope-tagged human Notch-1 (N1), a well-characterized POFUT1 substrate, was transiently overexpressed. Anti-FOT readily detected both full-length N1 and an extracellular domain (NECD) resulting from Furin cleavage in the Golgi, but not, as expected, an intracellular domain (NICD) (lane 3). Co-overexpression of LFNG, which caps the O-Fuc with a β3-linked GlcNAc, inhibited labeling (lane 2) confirming that anti-FOT reacted with O-Fuc. Similar findings were confirmed with anti-FLAG enriched material (lanes 4, 5). In contrast, anti-FOS failed to label overexpressed N1 (lane 8), even when the signal detection was dialed up to the point of labeling proteins non-specifically (e.g., see asterisk) or when N1 was enriched by immunoprecipitation (lane 10). This is consistent with the majority of O-Fuc being linked to Thr in N1, thus indicating specificity of anti-FOS toward FOS relative to FOT.

In a second approach, a truncated version of N1 consisting of EGF repeats 1-5, or a truncated version of human thrombospondin (TSP1) consisting of 3 TSRs, were purified from cell culture media of transiently transfected normal or glycosyltransferase mutant HEK293T cells (Fig. 6E). These constructs are targets of POFUT1 or POFUT2, respectively. As expected, anti-FOT (lane 11) but not anti-FOS detected truncated N1 (lane 15), and anti-FOT labeling depended on POFUT1 (lane 12), the enzyme that mediates addition of O-Fuc to EGF repeats. Purified truncated thrombospondin (TSP) was weakly detected by both anti-FOT and anti-FOS (lanes 13 and 17), and labeling was increased in the absence of the βGlc capping glycosyltransferase B3GLCT (lanes 14 and 18), confirming that reactivity was due to O-Fuc. In summary, endogenous levels of O-Fuc on these proteins was insufficient for detection by anti-FOS/T in whole cell extracts, but O-Fuc could be specifically detected when the proteins were expressed at high levels after purification. Significantly, thrombospondin repeat proteins modified by POFUT2 in *Toxoplasma* were also not detected in HFFs (Figs. 3,4), consistent with efficient capping with endogenous β3GlcT (14,47). Further studies are required to assess whether anti-FOS/T recognize FOS/T in the context of the disulfide stabilized folded domains or require denaturation as might occur during SDS-PAGE and western blotting. The findings show that O-Fuc recognition is blocked by capping sugars, and the comparisons indicate that anti-FOS preferentially recognizes FOS over FOT, suggesting anti-FOS does not tolerate the β-methyl substitution found in Thr.

## DISCUSSION

### Specificity of antisera for O-Fuc

This study provides a proof-of-principle that pan-specific anti-FOS and anti-FOT IgG antibodies can be generated by rabbits and used to localize O-Fuc proteins in cells and enrich them to define the O-fucome and map O-Fuc attachment sites. The antisera were generated against libraries of synthetic fucopeptides whose FOS or FOT flanking positions were occupied by amino acids highly represented in a study of O-Fuc proteins from *Toxoplasma* (5). These FOS-peptide and FOT-peptide libraries consistently induced high titer antisera which, after purification as described in Fig. 1A, titered out to beyond 10-30 ng/ml by ELISA, and had insignificant reactivity to the unmodified peptides.

Anti-FOS and anti-FOT were also specific for O-Fuc at the protein level. Based on western blot analysis of whole cell extracts of *Toxoplasma*, *Arabidopsis,* and *Dictyostelium*, multiple bands were detected with each (Figs. 3,5,6). At the illumination levels used to detect O-Fuc proteins, recognition was abolished in *spy*-KO cells (or reduced in *Arabidopsis* hypomorphs) demonstrating a dependence on this enzyme, which is a documented OFT in *Toxoplasma* (12). Similarly, no reactivity was observed in human fibroblasts, consistent with the absence of OFT in animals. Recognition was abolished by αMeFuc, confirming the presence of Fuc in the epitope recognized by the pAbs. Comparison of the western blot *M*_r_ profiles indicate distinct specificities for anti-FOS and anti-FOT presumably due to Ser or Thr being the attachment point for the Fuc residue. This possibility is corroborated by proteomic analysis of the captured *Dictyostelium* proteins (Table 2). Approximately 90% of the vegetative stage proteins that satisfied criteria for being classified as O-Fuc proteins were exclusive to the anti-FOS or anti-FOT pulldowns, and similar findings (80% exclusivity) were obtained at the slug stage. Since O-Fuc tends to be enriched in Ser-rich and Thr-rich regions, this correlation suggests that the anti-FOS and anti-FOT pAbs have considerable preference for FOS and FOT, respectively. O-Fuc proteins appear to be efficiently captured by anti-FOS/T based on the finding that >70% (or >83% at a lower threshold) O-Fuc proteins captured by AAL from a cytosolic preparation of *Dictyostelium* amoebae (7) were selectively captured by these pAbs. Additional proteins were detected only by anti-FOS/T, which indicated greater efficiency but also might be due to access to proteins that were not in the cytosolic preparation used for AAL. A direct comparison of AAL, anti-FOS and anti-FOT *M*_r_ profiles in *Toxoplasma* (Fig. 3D), where little other unmasked fucosylation occurs, suggests that at least the majority of AAL-reactive bands are detected by either anti-FOS or anti-FOT. Assessment of differences between AAL and anti-FOS/T is confounded by AAL-biotin also detecting intrinsically biotinylated proteins, and possible avidity differences between AAL and IgG, which are both bivalent, toward O-Fuc targets that vary in density and spacing.

**Table 2.**
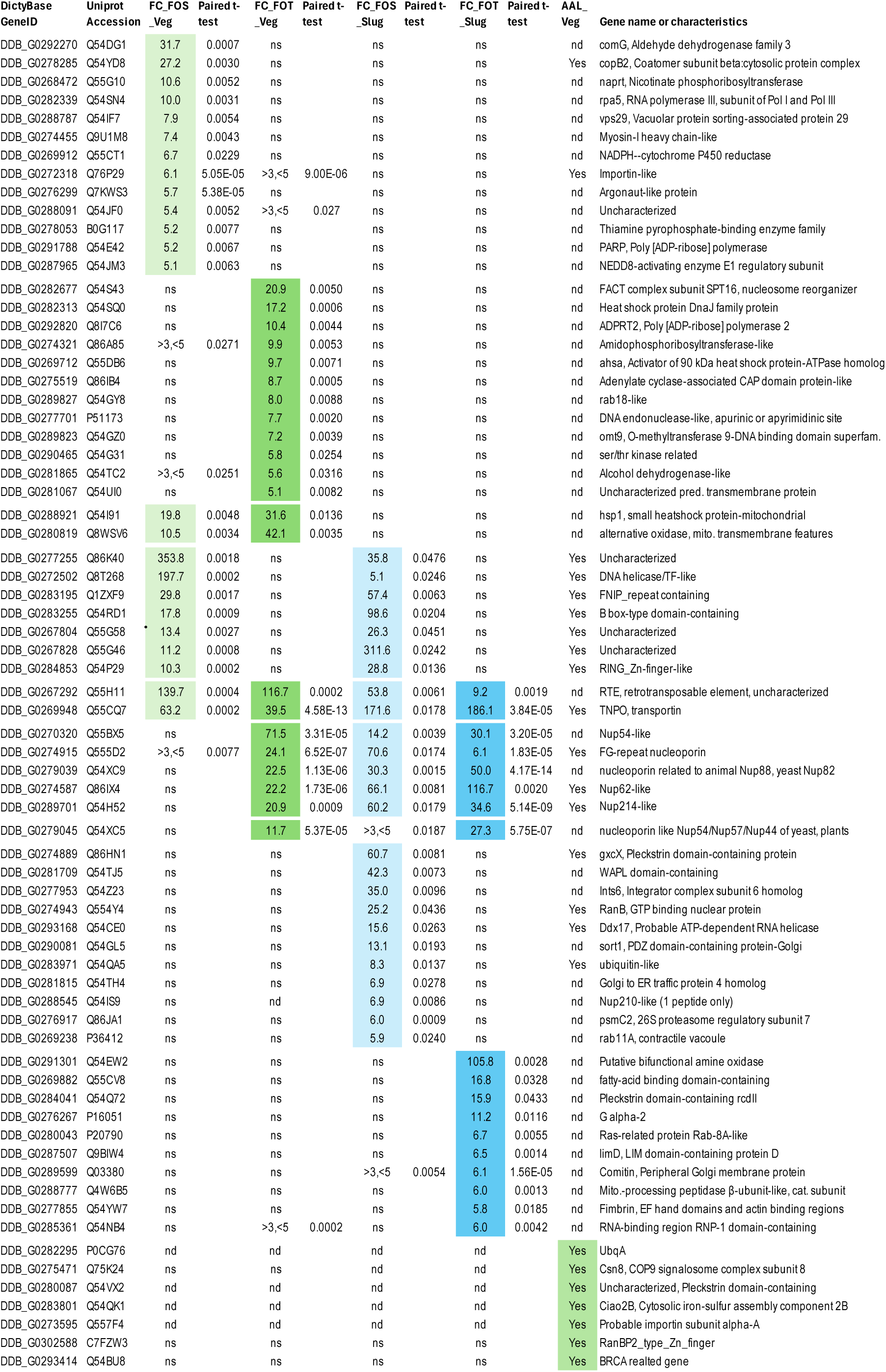
*Dictyostelium* O-Fuc protein candidates.

The specificity of the anti-FOS and anti-FOT Abs were also confirmed by immunofluorescence localization of fixed cells. Labeling was dominant in nuclei of *Toxoplasma*, *Acanthamoeba* and *Dictyostelium* and enriched in circumferential assemblies (Figs. 2,4,6). No clear differences were observed between anti-FOS and anti-FOT at this level of resolution. The considerable co-localization of O-Fuc with two *Dictyostelium* nuclear proteins (Figs. 4D,S3D) agrees with the finding of 7 FG-repeat nuclear pore proteins in AAL, anti-FOS, and anti-FOT pulldowns (Table 2). Minimal signal was detected outside of the nucleus despite the high level of other types of fucosylation detected with AAL in the secretory pathway, at the cell surface, and presumably in the endosome/lysosome system. The residual signal detected in the cytoplasm likely represents diffuse distributions of O-Fuc proteins. These findings are reminiscent of original immunofluorescence localization studies in *Toxoplasma* indicating a concentration of O-Fuc proteins in assemblies that subtended the nuclear envelope in register with nuclear pores (5). Nuclear enrichment of O-Fuc is consistent with the presence of OFT in the nucleus and the cytoplasm of *Toxoplasma* (12). Remarkably, the immunofluorescence findings parallel the original localization of O-GlcNAc in mammalian cells (49), which is mediated by the OFT-paralog OGT.

### Breadth of the Spy-dependent O-fucome

Proteomic analysis of pulldowns with anti-FOS and anti-FOT considerably expanded the O-fucome of *Dictyostelium* to 70 from the 24 detected with AAL (7). Criteria used for classifying O-Fuc proteins were that they are i) present in the proteome captured by anti-FOS or anti-FOT at a level at least 5-fold higher in wild-type compared to *spy*-KO cells, ii) detected at a peptide FDR of <1%, iii) and detected by ≥2 peptides at a protein FDR of <5%. With these criteria, very few peptides from proteins expected to reside in mitochondria or in the secretory or endolysosomal pathways were detected. Although the present study was not designed to confirm the presence of fucopeptides in the mass spectrometry analysis, 3 examples are reported (Figs. S4-6). The overall numbers are comparable to those found in the first documentation of an O-fucome as described in *Toxoplasma* using AAL-pulldowns, which indicated 67 O-Fuc proteins, 33 of which were directly confirmed on the basis of mapping O-Fuc sites on peptides by MS/MS (5). Both O-fucomes included proteins involved in DNA replication, transcription, nuclear transport, translation, the ubiquitin proteasome system, protein trafficking, and metabolism. Recently, the O-fucomes of two higher plants (17,18) based on AAL pulldowns from purified nucleocytoplasmic cell fractions yielded hundreds of proteins, many of which were confirmed by MS/MS experiments that mapped attachment sites and included proteins participating in a wide swath of intracellular functions (17). A key finding from these studies is that O-Fuc is also highly enriched in Ser- and Thr-rich sequences, often homopolymeric and disordered. The target sequences are more Ser/Thr-rich than targets of OGT in animals, though the O-GlcNAcome is partially enriched in disordered regions as well. The comparisons indicate a high degree of conservation of the biochemical aspects of O-fucosylation in the protists and plants where the Spy gene is found (7).

The diversity of O-Fuc proteins contrasts with their high degree of apparent enrichment in the vicinity of nuclear pores (Figs. 4D,S3D). It seems likely that the immunofluorescence observations are biased by a high concentration of the O-Fuc epitope in association with nuclear pores, such that the apparently selective binding is an avidity effect. This interpretation is borne out by findings in *Toxoplasma* that GPN-GTPase-1, which is stoichiometrically and highly O-fucosylated on a Ser-rich domain, is highly enriched in the cytoplasm (unpublished data).

### Detection of Spy-independent O-Fuc

Ser and Thr residues residing in certain disulfide-stabilized peptide contexts can be directly fucosylated by several so-called polypeptide fucosyltransferases (POFUTs) in the rER of animal cells and *Toxoplasma* (44–47,14), but no reactivity of anti-FOS or anti-FOT was observed in human fibroblasts or HEK293T cells (Figs. 2,3,6D,6E). However, reactivity was observed when human Notch-1 (N1) or thrombospondin was overexpressed; in these instances recognition depended on the appropriate POFUT and inhibited by overexpression of the appropriate glycosyltransferase that normally caps the O-Fuc. Furthermore, N1, which is largely modified at Thr residues, was not recognized by anti-FOS, supporting amino acid specific recognition of O-Fuc. Anti-FOS/T also captured a class of spore coat proteins in *Dictyostelium* that have previously been documented to be modified in Ser/Thr-rich mucin-like domains with O-Fuc, and enabled detection of an O-Fuc peptide (Fig. S6). Like Notch and TSP, these proteins are modified in the secretory pathway but by a currently unknown fucosyltransferase. The detection in both wild-type and *spy*-KO strains confirms lack of dependence on the Spy OFT. Notably, four other spore coat proteins with Ser/Thr-rich mucin-like domains were present in the pull-downs, whereas an equally abundant spore coat protein, SP60 which lacks mucin-like domains (49), was barely represented. These findings supported the broad ability of anti-FOS and anti-FOT to detect FOS and FOT on any protein, including the minimal level of uncapped FOS and FOT in normal cultured cells. In addition, anti-FOS was specific for FOS relative to FOT at the protein level. However, it is unclear whether anti-FOS and anti-FOT can detect uncapped O-Fuc in the context of their disulfide-stabilized folded domains prior to denaturation.

### Regulation of O-Fuc

A previous analysis showed that O-Fuc disappears after *Toxoplasma* differentiates into oocysts and reappears in tachyzoites (5). The basis for this regulation is unclear, but an α-fucosidase that might remove O-Fuc has not been identified. In *Dictyostelium*, a dramatic increase in O-Fuc occurred as cells differentiated from the growth stage to multicellular organization and differentiation into stalk and spore cells in fruiting bodies (Figs. 5D,E). It is unclear whether this involves a change in the specificity or general activity of OFT, or a change in the repertoire of proteins available to be O-fucosylated. Given the potential absence of a mechanism to remove O-Fuc, O-Fuc proteins might be simply removed by proteome remodeling or dilution by proliferation. Though biochemical functions of O-Fuc are not known, other monosaccharide modifications have been shown to influence local peptide and protein conformations (50). The breadth of the O-fucome in protists invites speculation that the roles of O-fucosylation may have parallels with the roles of animal O-GlcNAc in responding to stress and nutritional changes.

## MATERIALS AND METHODS

### Antisera production

Four dodecapeptides were synthesized by AnaSpec (Fremont, CA) using FMOC chemistry. The glycoamino acid conjugates FMOC-Ser-Fuc and FMOC-Thr-Fuc were purchased from Sussex Research (San Diego, CA). The sequences were CXXXXX-Y-XXXXX, where Y was, Ser, Thr, Fuc-O-Ser (FOS), or Fuc-O-Thr (FOT). X indicates a mixture of 10 amino acids in the following molar percentages based on flanking sequences of fucopeptides from *T. gondii* (5): Ser 30%; Gly 15%; Ala 15%; Thr 10%; Phe 5%; Pro 5%; Val 5%; Leu 5%; Arg 5%; Gln 5%. The fucopeptides were incorporated via their Cys-residues, by GlycoScientific LLC, into a proprietary three-component immunogen composed of the fucopeptides covalently attached to a helper T-cell (CD4+) epitope sequence and a Toll-like receptor-2 agonist as a built-in adjuvant (51,52). This approach has been successful in the generation of glycopeptide specific rabbit antisera (53,54) and mouse mAbs (55), where rabbits have been more robust than inbred mice for generating glycopeptide responses (56). After coupling to keyhole limpet hemacyanin, the FOS and FOT immunogens were separately used to immunize and boost two pairs of female New Zealand white rabbits. For ELISA screening, peptides were dissolved in water and conjugated to bovine serum albumin (BSA) using Thermo Scientific™ Imject™ Maleimide-Activated BSA (catalogue #PI77667). The peptide/BSA conjugates were adsorbed to microwell plates, incubated with a dilution series of antisera, and antibody binding quantified using alkaline phosphatase conjugated AffiniPure Goat Anti-Rabbit IgG from Jackson ImmunoResearch (#111-055-046). For negative absorption or positive affinity purification as illustrated in Fig. 1A, peptides were bound to Thermo Scientific™ UltraLink™ Iodoacetyl resin (#53155) beads per the Manufacturer’s protocol. Agnostic and affinity anti-FOS/T pAbs were eluted with 0.1 M glycine-HCl (pH 2.5), neutralized, concentrated to 1.0 mg/ml and amended with 0.07% Tween-20, and dialyzed against 0.07% Tween-20, 0.05% Proclin in PBS (pH 7.4). Each rabbit generated high titers of anti-fucopeptide pAbs based on ELISA assays. The data in this report were derived primarily affinity-purified Abs from one rabbit from each set, and confirmatory findings from other rabbits are not detailed.

### *Toxoplasma* tachyzoite cell culture

RHΔ*ku80*Δ*hxgprt* (RHΔΔ) and RHΔΔSPYΔ type 1 *Toxoplasma* strains (Table 1) were propagated in human telomerase reverse transcriptase immortalized foreskin fibroblasts (hTERTs) (57) and transferred to primary human foreskin fibroblasts (HFFs) for experimental procedures (58). Cultures were maintained at 37°C, 5% CO_2_, in Dulbecco’s Modified Eagle Medium (DMEM), supplemented with 1% (v/v) fetal bovine serum (FBS), 200 mM glutamine, 100 units/ml penicillin, and 100 μg/ml streptomycin. Parasites were harvested from HFF hosts grown in T25 or T175 flasks inoculated with spontaneously lysed extracellular tachyzoites and grown for 2 days, or until parasites began to spontaneously lyse out. For western blotting, parasites were released from scraped host monolayers by four passages through a 27-gauge syringe needle, resuspended in DMEM, and centrifuged at 2000 × *g* for 8 min at 22°C. Parasites were resuspended in PBS and centrifuged at 1800 × *g* for 10 min, and the pellet frozen at –80°C or immediately solubilized for SDS-PAGE.

### Immunofluorescence analysis of *Toxoplasma gondii*

HFF monolayers prepared on 12 mm diameter glass coverslips in 24-well plates were infected with tachyzoites and later fixed in cold methanol for 2 min, washed thrice with PBS (Corning 21-040-CV), and incubated in Blocking Buffer (5% (v/v) FBS, 5% (v/v) normal goat serum in PBS) for an additional 10 min. Coverslips were exposed to the following primary antibodies diluted in Blocking Buffer: affinity purified rabbit anti-FOS (1 µg/ml), rabbit anti-FOT (0.3 µg/ml), mouse anti-SAG1 (1:5000, MyBioSource), mouse anti-PLP6 (1:10,000, Millipore Sigma), and/or biotinylated *Aleuria Aurantia* Lectin (AAL) premixed with Alexa Flour 594 streptavidin (1:250, Vector Labs), for 1 h at 22°C. Goat anti-mouse and anti-rabbit secondary antibodies (Invitrogen) conjugated to Alexa Fluor 488 or 594 were applied at a 1:10,000 dilution in Blocking Buffer for 1 h at 22°C. After 2 final washes with PBS and once quickly with water, coverslips were mounted with Prolong Antifade Diamond with DAPI (Molecular Probes) on glass slides and allowed to dry overnight at 4°C.

Samples were imaged with a Zeiss ELYRA S1 Super Resolution-Structured Illumination Microscope (SR-SIM) and processed with the supporting SR-SIM ZEN 2011 software. Images were acquired with 63x/1.4 NA or 100x/1.4 NA oil immersion objectives with 0.11 μm z sections. Pearson’s correlation coefficients were calculated from super-resolution images utilizing the Coloc 2 software from FIJI/Image J (59,60).

### Immunofluorescence analysis of *Dictyostelium discoideum*

∼10^6^ vegetative (growth stage) amoebae were seeded onto 12-mm diameter glass coverslips coated with poly-L-lysine (1 μg/ml, Sigma Aldrich) and allowed to adhere for 30 min at 22°C. Adherent amoeba were washed thrice with PBS and fixed with ice-cold methanol for 30 min at 4°C. Samples were washed thrice with PBS and blocked with 5% (w/v) BSA in PBS for 10 min. Coverslips were exposed to the following primary antibodies diluted in 5% BSA in PBS: rabbit anti-FOS (2 µg/ml), rabbit anti-FOT (0.6 µg/ml), mouse anti-actin (1:5000, Sigma Aldrich), mouse anti-PLP6 (1:5000), and/or biotinylated AAL (1:250, Vector Labs) pre-bound to Alexa Flour 594 streptavidin for 1 h at 22°C. Secondary antibodies, coverslip mounting and imaging were as described above.

### Immunofluorescence analysis of *Acanthamoeba castellanii*

Trophozoites of the *Acanthamoeba castellanii* Neff strain were grown in axenic culture at 30°C in T75 tissue culture flasks as described (62). Ten million trophozoites were fixed in 4% paraformaldehyde for 15 min at 22°C, permeabilized with 0.1% Triton X-100 for 10 min, blocked with 1% (w/v) BSA for 1 h, and incubated with 1:500 fluorescein-AAL (Vector Laboratories, Newark, CA), anti-FOS rabbit antibodies, or anti-FOT rabbit antibodies for 1 h at 22°C, followed by washing and incubation with anti-rabbit secondary antibodies conjugated to Alexa Fluor 488 (1:500). Confocal microscopy of DAPI-labeled trophozoites was performed as described (62).

### *Arabidopsis* sample preparation for western blot analysis

Whole seedlings (Table 1) frozen in liquid nitrogen were ground to a fine powder utilizing a wooden stick and transferred to 1.5 ml polypropylene tubes. The powder was extracted twice with 1 ml of acetone at −20°C for 30 min and recovered by centrifugation at 17000 × *g* for 2 min. The process was repeated using 500 µl of acetone and incubation for 5 min at 22°C. Acetone was removed, samples were dried by vacuum centrifugation, and resuspended in 3.5 ml/g fresh weight of tissue with 50 mM Tris-HCl (pH 8.0), 1% SDS. Solubilized samples were heated at 65°C for 15 min with occasional vortexing, and boiled for an additional 5 min. Protein concentration was determined using Bicinchoninic acid according to the manufacturer’s instructions (BCA Kit, Thermo Fisher Scientific).

### HEK293 samples

HEK293T cells (ATCC) were cultured in 5% CO_2_ at 37°C in DMEM high glucose media with 10% bovine calf serum. *B3GLCT1*-null (35) and *POFUT1*-null (34) HEK293T cells were described previously. The following plasmids were used: pSecTag-mouse LFNG with myc and 6×His tag (63), pSecTag-mouse N1 EGF1-5 with MYC and 6×His tag (64), pSecTag-human TSP1 TSR1-3 with MYC and 6×His tag (65), pcDNA5-FRT-TO-human N1 with N-terminal (extracellular) FLAG and C-terminal (intracellular) myc tag (66).

Transfection and protein purification methods were described previously (67). Briefly, for N1 EGF1-5 or TSP1 TSR1-3 transfection, 5 μg of pSecTag-N1 EGF1-5 or TSP1 TSR1-3 with 30 μg of PEI reagent were used for each of three 10 cm dishes of HEK293T cells. After 4-6 h incubation, cells were washed twice with 10 ml of warm PBS and incubated with 10 ml of OPTIMEM for 72 h. Secreted proteins were purified from culture media using 100 μl of 50% slurry of Ni-NTA (Qiagen). After overnight rotation at 4°C, Ni-NTA was washed with 1 ml of TBS-8 (100 mM NaCl, 50 mM Tris-HCl, pH pH 8.0) with 10 mM imidazole and 0.5 M NaCl, and eluted with 250 μl of 100 mM imidazole, TBS-8. For FLAG-N1-myc, 10 cm dishes of HEK293T cells were each transfected with 3 μg of pcDNA5-FRT-TO-human N1 and 1.5 μg of pSecTag-LFNG with 30 μg of PEI reagent in DMEM and incubated for 48 h. Cells were washed with TBS-8 three times and lysed with 400 μl of 1% NP40/TBS-8 with protease inhibitor (Thermo). The cell lysate was centrifuged at 16,260 × *g* for 10 min at 4°C and used for FLAG IP. The lysate was incubated with 60 μl of a 50% slurry of anti-FLAG beads (Sigma) on a tilting rotator at 4°C for 6 h. Beads were collected using a magnet and thrice washed with 1% NP40, TBS-8 and thrice with TBS-8. N1 was eluted from the beads by boiling with SDS sample buffer containing 2-mercaptoethanol.

### SDS-PAGE and western blotting

*Toxoplasma* pellets or host cells were solubilized in Laemmli Sample Buffer containing 50 mM dithiothreitol, heated at 100°C for 3 min, and loaded at the equivalent of 5×10^6^ parasites/lane (unless otherwise indicated) on a 1-mm thick 4%-12% Bis/Tris NuPage polyacrylamide SDS gel (Invitrogen). Axenic strains of *Dictyostelium discoideum* (Table 1) were grown and subjected to starvation induced development to form fruiting bodies as described (7). *Dictyostelium* vegetative and developing cells were similarly prepared and loaded at 2.5×10^5^ or 5×10^5^ cells/lane respectively. *Arabidopsis* seedling extracts were loaded at 50 µg total protein/lane. Samples were electrophoresed using MOPS running buffer, and gels were either stained with Coomassie blue or transferred to a nitrocellulose membrane using an iBlot 2 (Invitrogen) dry blotting system, as described (58,7). Blots were blocked with 5% (w/v) nonfat dry milk in Tris-buffered saline (TBS) (100 mM NaCl, 50 mM Tris-HCl, pH 7.5), followed by overnight incubation with anti-FOS (2 µg/ml), anti-FOT (0.6 µg/ml), or biotin-AAL (1 µg/ml) diluted in the same blocking solution. After washing thrice with TBS-7.5, blots were probed with Alexa Fluor-680 goat-anti rabbit IgG (1:20,000) secondary Abs, or Alexa Fluor-680 streptavidin (1:10,000). Blots were scanned for fluorescence intensities with a LiCor Odyssey CLx scanner (Li-COR Biosciences, Lincoln, USA).

For HEK293T extracts, IP samples or cell pellets resuspended in SDS sample buffer containing 2-mercaptoethanol were subjected to SDS-PAGE using a 4%-15% pre-cast gradient gel (BioRad), and transferred to a nitrocellulose membrane (BioRad) in transfer buffer containing 10% methanol at 22°C for 60 min. Membranes were blocked with TBS-7.5 containing 0.1% Tween-20 (TBST) and 5% BSA at 22°C for 30 min, and incubated overnight at 4°C with anti-myc (1/1000, 9E7), anti-FOT (0.3 μg/ml), and/or anti-FOS (1 μg/ml). Membranes were washed thrice with TBST for 15 min and incubated at 22°C for 1 h with anti-mouse IgG IRDye 680 (1:5000, Li-COR #925-68970) and anti-rabbit IgG IRDye 800 (1:5000, Li-COR #926-32211). Membranes were washed thrice with TBST and TBS for 15 min, and scanned as above.

### Immunoprecipitation of *Dictyostelium* extracts

O-Fuc proteins were captured using a pulldown approach as described (7 et al. 2023). Briefly, cells (2-3×10^5^/µl) were solubilized in Lysis Buffer (0.15 M NaCl, 20 mM Tris-HCl, pH 7.5, 0.5% (w/v) CHAPS, 1 mM DTT, 10 μg/ml leupeptin, and 10 μg/ml aprotinin) utilizing a probe sonicator on ice, and centrifuged at 21,000 × *g* for 15 min to remove the insoluble material yielding an S21 fraction. 1.0-1.2 mg of each affinity purified pAb anti-FOS or -FOT were conjugated and covalently crosslinked (68) to 100 μl of protein A/G magnetic agarose beads (Pierce, 78610). To enrich for O-Fuc proteins, 5×10^7^ cell equivalents of S21 extracts were used to resuspend 2.5 µl packed volume of the anti-FOS/T magnetic beads. After rotation for 1 h at 4°C, the beads were collected magnetically, rinsed 3× in detergent-free buffer (10 mM Tris-HCl, pH 7.5, 50 mM NaCl), and twice more in 100 µl of 50 mM NaCl. For western blot and mass spectrometric analysis, beads were incubated for 15 min and eluted with 100 µl of 0.2 M αMeFuc, 8 M urea at 22°C. Samples were then reduced, alkylated, and converted to peptides with endo LysC and trypsin; peptides were then captured on C18 Zip-tips and released for MS analysis, all essentially as described (69).

### Mass spectrometry of peptides

Peptides were separated on a C18 nano-column (Thermo Acclaim PepMap 100 C18 series) in a 5% to 40% acetonitrile 75 min linear gradient using an Ultimate 3000 nano-HPLC, and directly infused into a Q-Exactive Plus Orbitrap Mass Spectrometer (Thermo Fisher). Peptides were fragmented in the C-trap via stepped HCD collision energies of 17, 27 and 37 for MS/MS analysis using a Top 20 method and a 10 sec fragmentation exclusion window. Raw files were processed in Proteome Discoverer 2.5 using the *D. discoideum* protein database as described (69), with the following modifications: dynamic oxidation of Met, dynamic loss of N terminal Met with or without acetylation of N terminal Met, dynamic addition of ≤2 dHex (+146.058 Da), and static carbamidomethylation of Cys. From 3 biological replicates of both vegetative and slug cells, each including three technical replicates, Proteome Discoverer 2.5 calculated protein abundances from ≥2 peptides detected at FDR<0.01 at high (FDR<0.01) and medium confidence (FDR<0.05). Using MetaboAnalyst 5.0 (70). proteins enriched >5-fold with a p<0.05 in w/t vs. *spy*-KO samples were assigned as O-Fuc proteins. Data are deposited in the ProteomeXchange Consortium via the PRIDE (71) partner repository with the dataset identifiers PXD PXD056853, PXD056857, PXD056903, PXD056866 and PXD057018.

## Supporting information

Revised Supplement file

## ABBREVIATIONS

AAL: *Aleuria Aurantia* Lectin
Ab: antibody
αMeFuc: Fuc-1-Me
anti-FOS: affinity-purified anti-FOS peptide library antisera
anti-FOT: affinity-purified anti-FOT peptide library antisera
ATCC: American Type Culture Collection
BSA: bovine serum albumin
co-IP: co-immunoprecipitation
FOS: fucose-O-Ser
FOT: fucose-O-Thr
Fuc: α-L-fucose
DAPI: 4’,6-diamidino-2-phenylindole
HFF: human foreskin fibroblast
hTERTs: human telomerase reverse transcriptase immortalized foreskin fibroblasts
IP: immunoprecipitation
KI: knock-in or addition of genetic coding information
KO: knock-out or removal of genetic material generating a null allele
PBS: phosphate buffered saline (without divalent cations)
SR-SIM: Super-Resolution Structured Illumination Microscopy
wild-type: refers to the normal parental strain used in this study

## ACKNOWLEDGMENTS

We thank Muthugapatti K. Kandasamy for his assistance with the light microscopy.

## DATA AVAILABILITY

The MS proteomics data, which are indexed in Table S2, are deposited in the ProteomeXchange Consortium via the PRIDE (71) partner repository with the dataset identifiers PXD PXD056853, PXD056857, PXD056903, PXD056866 and PXD057018.

## AUTHOR CONTRIBUTIONS

M.T., E.G.-P., M.G., M.P., and N.O. performed the experiments. M.T., R.O., J.S. and C.M.W. conceived the project and experimental approaches. M.T wrote the first draft of the manuscript, C.M.W. was the primary editor, and all authors reviewed and approved of the final submission.

## ADDITIONAL FILES

### Supplemental Material

This article contains supporting information, with two tables and six figures.

