## Supplementary material for "Novel antibodies detect nucleocytoplasmic O-fucose in protist pathogens, cellular slime molds, and plants": Revised Supplement file

**Table of Contents**

**Figure S1.** Screening of affinity purified anti-FOT and agnostic Abs against parasite infected HFFs using super-resolution microscopy.

**Figure S2**. Immunofluorescence analysis of co-labeling of anti-FOS or anti-FOT with anti-PLP6 or AAL in *Toxoplasma* infected HFFs.

**Figure S3**. Immunofluorescence analysis using anti-FOS/T on vegetative stage (growing) *Dictyostelium*.

**Figure S4.** Mass spectrometric confirmation of 2 fucopeptides from a putative nucleoporin.

**Figure S5.** Mass spectrometric confirmation of a fucopeptide in a putative helicase/transcription factor.

**Figure S6.** Mass spectrometric confirmation of fucopeptides from the spore coat protein SP70.

**Table S1.** Proteins found at increased levels in anti-FOS/T pulldowns of *spy*-KO cells.

**Table S2.** List of proteomics files available at the ProteomeXchange Consortium via the PRIDE partner repository.

**Figure S1.** Screening of affinity purified anti-FOT and agnostic Abs against parasite infected HFFs using super-resolution microscopy. (A) HFFs containing wild-type RHΔΔ (upper row) or TgSPYΔ (middle row) parasites were probed with rabbit anti-FOT (0.3 µg/ml) and murine anti-SAG1 (1:1000) followed by Alexafluor-488 goat anti-rabbit IgG and Alexafluor-594 goat anti-mouse IgG to localize O-Fuc and outline the parasites, respectively. DAPI (blue) was used to visualize nuclei. Lower row: RHΔΔ-infected HFFs were probed with anti-FOT in the presence of 0.2 M alpha-methyl fucose (αMeFuc). Samples were imaged using a Zeiss ELRYA S1 microscope and processed with SR-SIM, and maximum projection images are shown. Scale bars: 5 µm. See Fig. 2 for corresponding probing with anti-FOS. (B, C) Agnostic pAB libraries created from affinity purification of anti-FOT (Thr) or anti-FOS (Ser) (see Fig. 1A) showed no distinguishable labeling in the presence or absence of SPY.

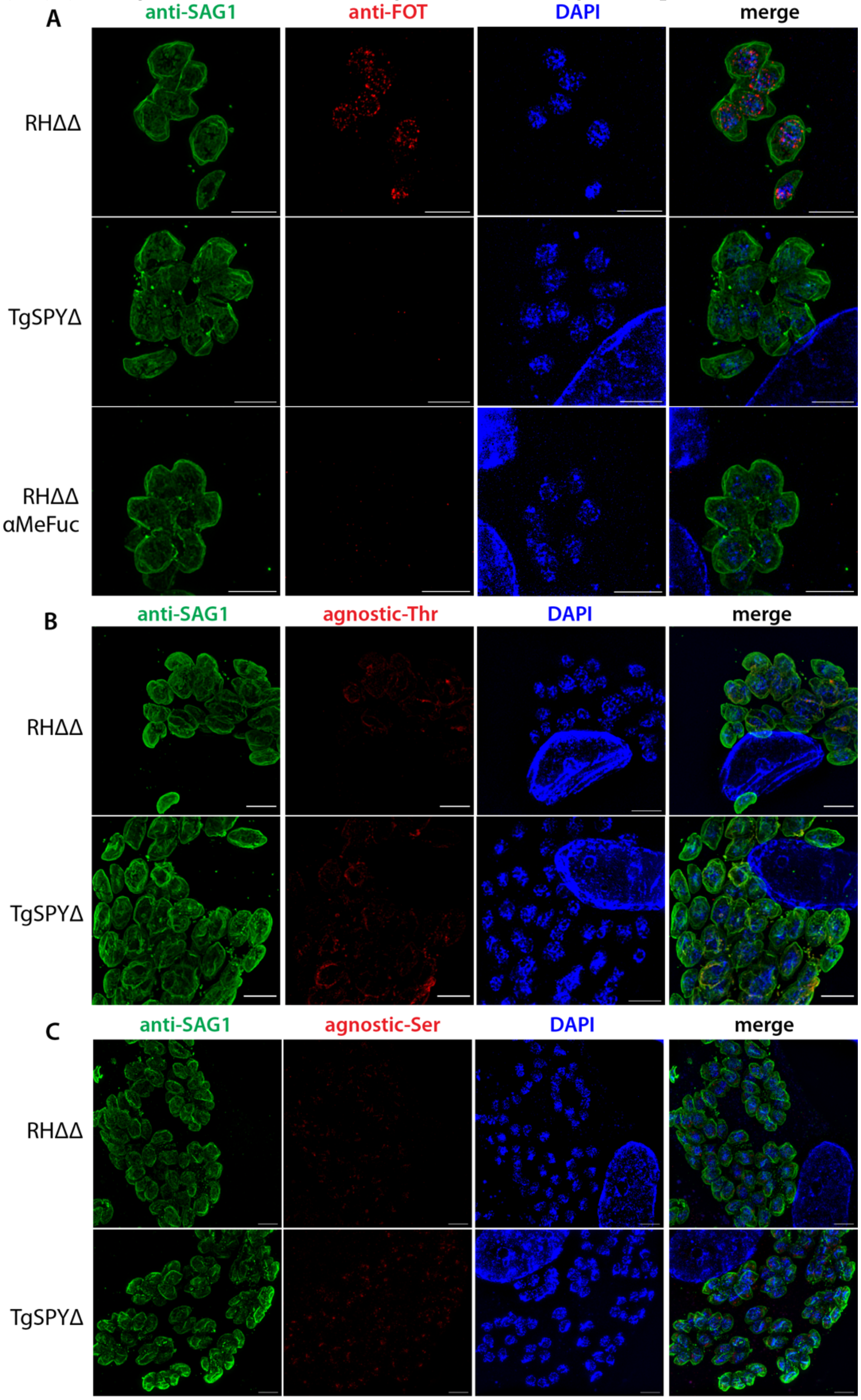

**Fig. S2**. Immunofluorescence analysis of co-labeling of anti-FOS or anti-FOT with anti-PLP6 or AAL in *Toxoplasma* infected HFFs. (A) HFFs containing wild-type RHΔΔ were probed with affinity purified rabbit anti-FOS (1 μg/ml) or anti-FOT (0.3 μg/ml) and mouse anti-PLP6 followed by Alexa Fluor-488 goat anti-rabbit IgG and Alexa Fluor-594 goat anti-mouse IgG. (B) Same as above, except that samples were probed with anti-FOS or anti-FOT and AAL. Samples were imaged as in Fig. S1. Scale bars: 5 μm.

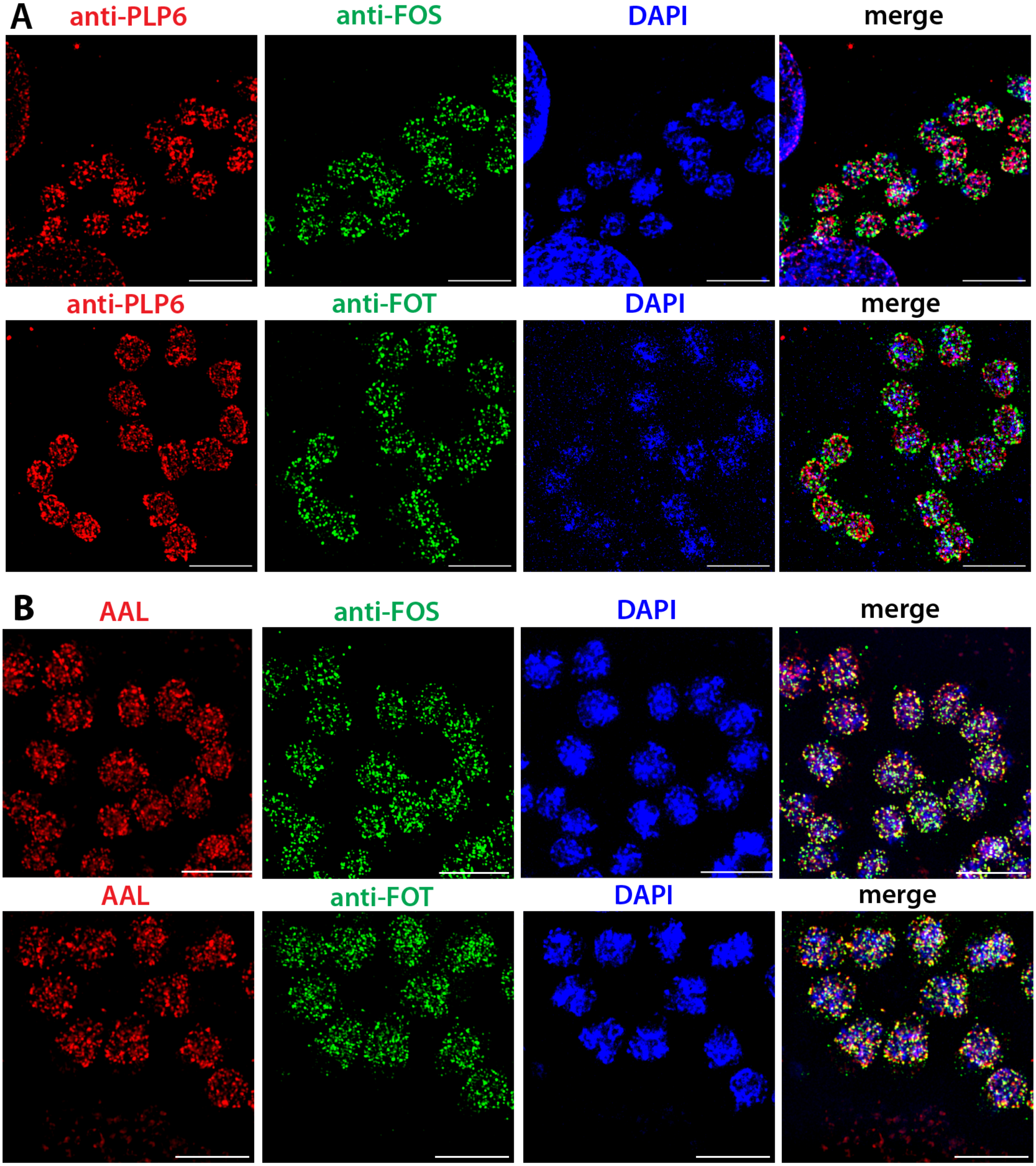

**Figure S3**. Immunofluorescence analysis using anti-FOS/T on vegetative stage (growing) *Dictyostelium*. Growth stage amoebae were allowed to attach to cover slips and then fixed and permeabilized with MeOH. (A) wild-type (w/t, strain Ax3) amoebae were probed with anti-FOT and AAL-biotin, followed by Alexa Fluor-488 goat anti-rabbit IgG and Alexa Fluor-594 streptavidin. DAPI (blue) was used to visualize nuclei. Maximum projection images are shown. (B) Similarly, w/t, Dd*spy*⁻ (KO), and Dd*gft*⁻ amoebae were probed with anti-FOT (0.6 µg/ml) and anti-actin (1:250) followed by Alexa Fluor-488 goat anti-rabbit IgG (1:250) and Alexa Fluor-594 goat anti-mouse IgG (1:250). (C) Same as panel A, except that amoebae were probed with anti-FOS and anti-PLP6 (1:5000), followed by Alexa Fluor-488 goat anti-rabbit IgG and Alexa Fluor-594 goat anti-mouse IgG. (D) Comparison of anti-FOT with nuclear envelope proteins. Amoebae whose Nup62, Nup210, or Src1 genomic loci were C-terminally tagged with mNeon were probed with anti-FOT followed by Alexa Fluor-594 goat anti-rabbit IgG, and co-imaged with intrinsic mNeon fluorescence and DAPI. Single z-slice images are shown. Scale bars: 5 µm.

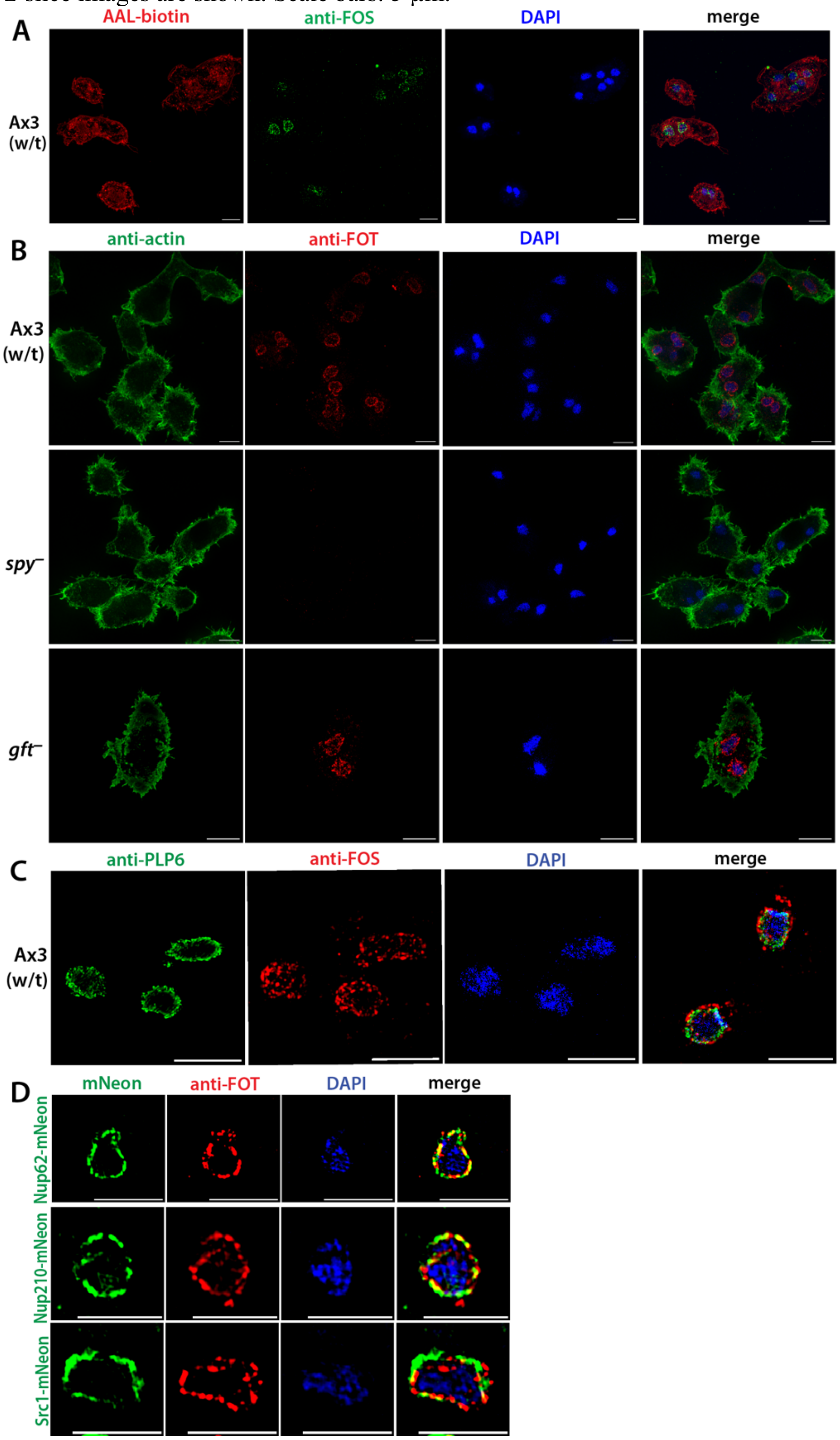

**Figure S4.** Mass spectrometric confirmation of 2 fucopeptides from a putative nucleoporin (dictyBase DDB_G0274915, Uniprot Q55D2). Peptides from an anti-FOS pulldown from wild-type *Dictyostelium* cells were profiled by nLC on a C18 column and introduced into an Orbitrap mass spectrometer operated in positive ion mode. (A) From vegetative cells, a triply-protonated ion group (a) was detected with a neutral loss of 146, corresponding to a fucose residue. The most abundant isomer (circled in green) was selected for secondary collision (b,c). Calculated and observed *m/z* values for the mono-fucosylated peptide are shown in (a) together with the peptide sequence and attachment site range inferred from analysis of b- and y-ions in panels b and c. Ions detected with loss of 146 are mapped with green arrows.

Q555D2, FG-repeat nucleoporin -> fucopeptide-1
AX3, anti-FOS, vegetative

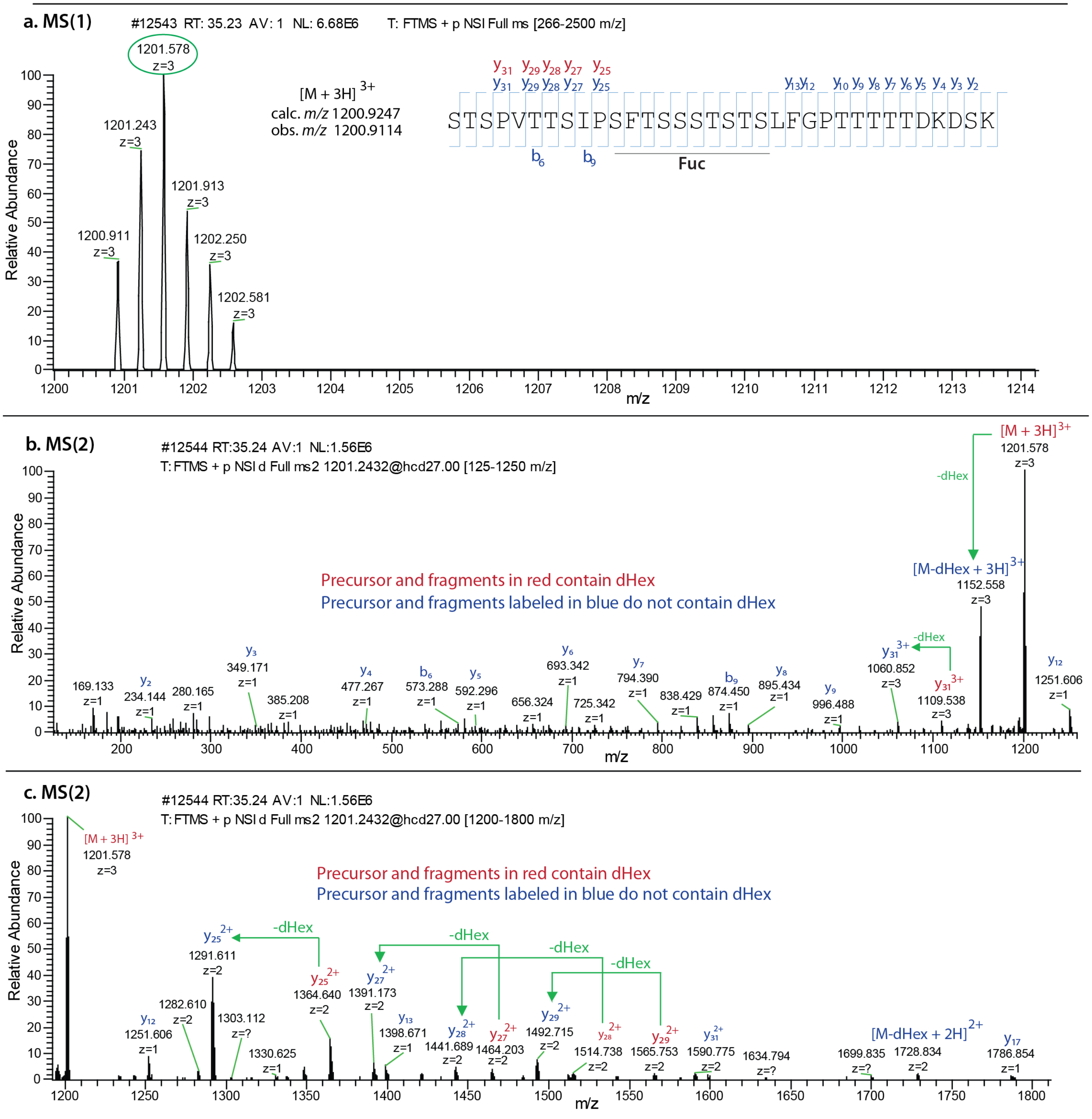

(B) Similarly for a second fucopeptide from the nucleoporin from slug cells. (a) A doubly-protonated ion that was also detected with a neutral loss of 146 was selected for secondary collision. (b) Zoom-in on the parent ion of the MS(2) profile showing the isotopic composition. (c) Zoom-out of the ion profile from which the sequence represented in panel A was deduced. The profile also shows that neutral loss of a fucose residue from the parent ion and y12 and y15 peptides, permitting partial mapping of the fucose to the Ser/Thr rich region of the peptide as indicated.

Q555D2, FG-repeat nucleoporin -> fucopeptide-2
AX3, anti-FOS, slug

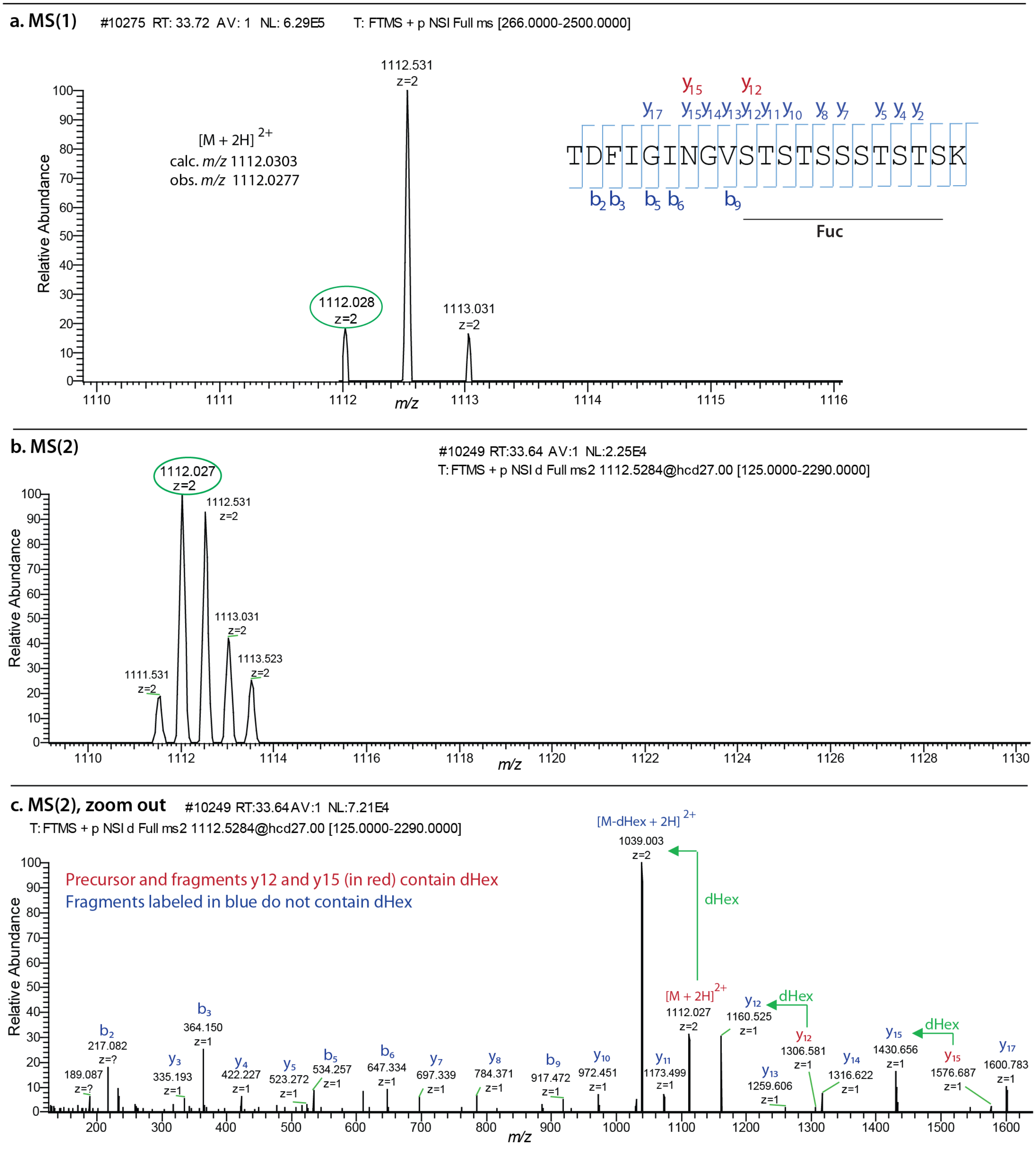

**Figure S5.** Mass spectrometric confirmation of a fucopeptide in a putative helicase/transcription factor (dictyBase DDB_G0272502, Uniprot Q8T268). Peptides from an anti-FOS pulldown from vegetative wild-type *Dictyostelium* cells were profiled by nLC on a C18 column and introduced into an Orbitrap mass spectrometer operated in positive ion mode. (A) An ion that was also detected with a neutral loss of 146, corresponding to a fucose residue, was selected for secondary collision. (B) MS(2) of the monoisotopic triply-charged parent ion yielded a series of fragment ions allowing assignment of its sequence as presented in panel A. The presence of a deoxyHex on the y25 fragment ion limits its possible location to the region indicated.

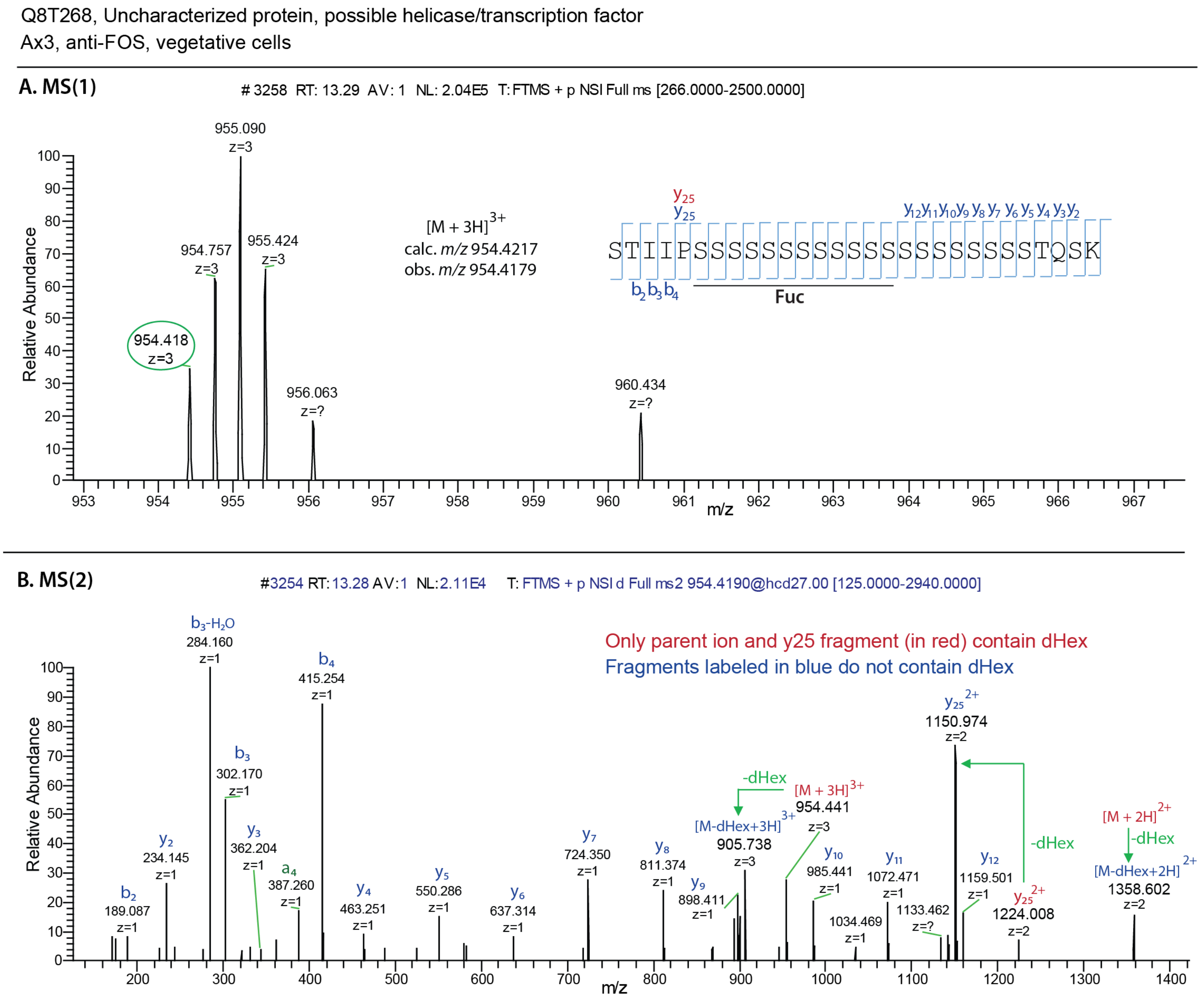

**Figure S6.** Mass spectrometric confirmation of fucopeptides from the spore coat protein SP70 (dictyBase DDB_G0276761, CotB, Uniprot P15269). Peptides from an anti-FOT pulldown from wild-type *Dictyostelium* slug cells were profiled by nLC on a C18 column and introduced into an Orbitrap mass spectrometer operated in positive ion mode. (A) Summary of findings. (B) An ion that was also detected with a neutral loss of 292, corresponding to 2 fucose residues, was selected for secondary collision. (C) MS(2) of the monoisotopic doubly-charged parent ion yielded a series of fragment ions allowing assignment of its sequence as presented in panel A. The presence of a deoxyHex on the y22 fragment ion limits its possible location to the region indicated. (D) Another ion that was also detected with a neutral loss of 146, corresponding to a single fucose residue, was selected for secondary collision. (E) Same as panel D.

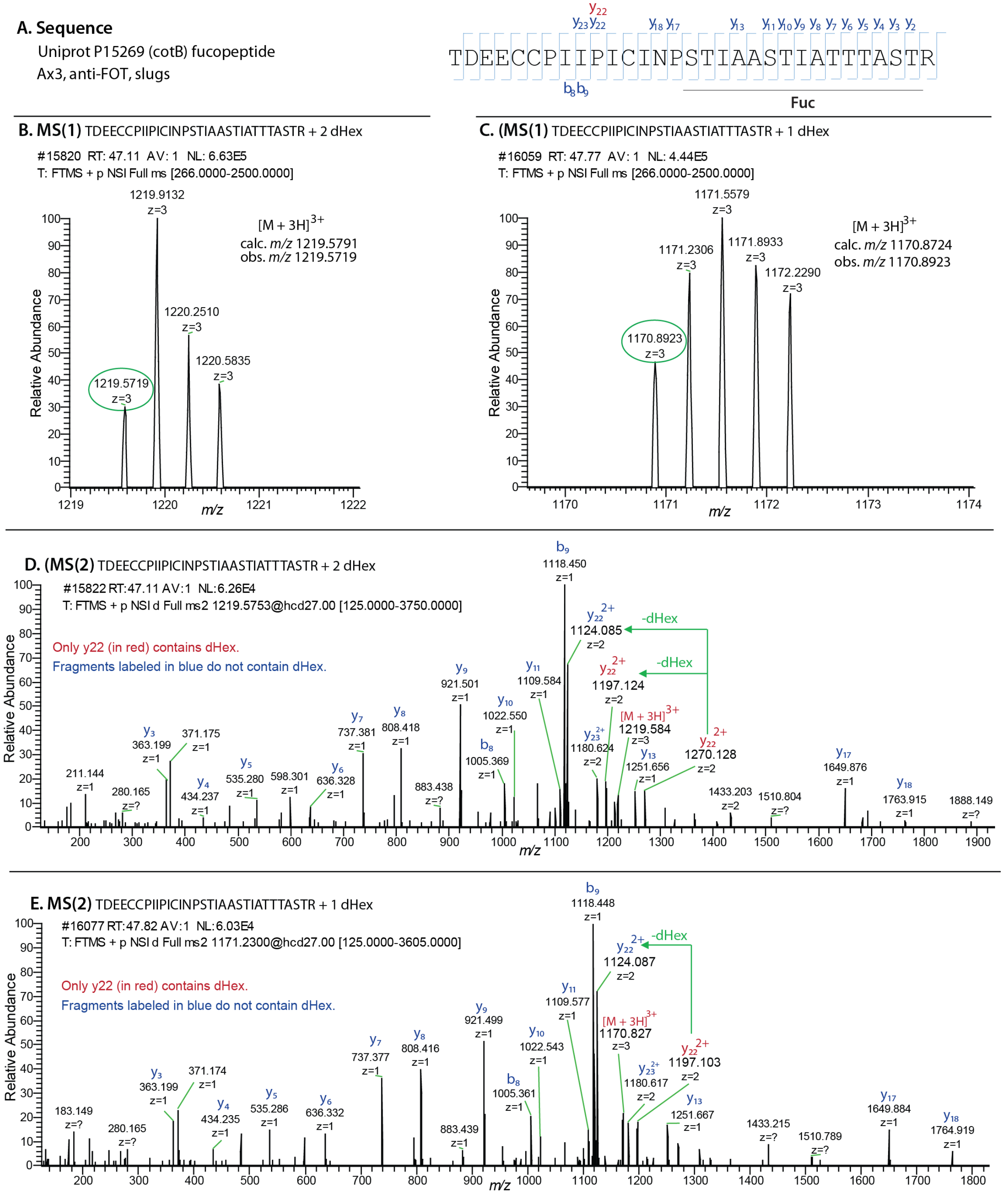

**Table S1.** Proteins found at increased levels in anti-FOS/T pulldowns of *spy*-KO cells. See volcano plots in Figs. 5E-H and associated text. Formatted is as in Table 2 of the main text. Columns represent fold-change (FC) for abundance based on spectral counting in wild-type vs. *spy*-KO cells using anti-FOS or anti-FOT in vegetative or slug stage cells as indicated. Significances of the fold-changes over the replicates are shown in the adjacent columns. ns, not significantly enriched in both anti-FOS and anti-FOT trials for the indicated stage; nd, not detected.

| Accession | DictyBase GeneID | Uniprot Accession | FC FOS Veg | Paired t-test | FC FOT Veg | Paired t-test | FC FOS Slug | Paired t-test | FC FOT Slug | Paired t-test | Short name/description |
|---|---|---|---|---|---|---|---|---|---|---|---|
| DDB0216434 | DDB_G0290331 | Q54G78 | 0.06 | 0.003 | 0.24 | 0.024 | nd | | nd | | ssrp1, FACT complex subunit SSRP1 |
| DDB0191196 | DDB_G0267456 | P54653 | 0.04 | 0.009 | 0.12 | 0.017 | ns | | ns | | cbp2, calcium-binding protein 2 |
| DDB0305759 | DDB_G0283653 | Q54QT0 | 0.07 | 5.14E -07 | 0.15 | 0.014 | nd | | nd | | Tyrosinase copper-binding domain protein |
| DDB0238141 | DDB_G0270212 | Q58A40 | 0.15 | 0.000 | 0.17 | 2.85 E-05 | ns | | ns | | Galactose-binding domain-containing protein |
| DDB0216703 | DDB_G0270952 | Q55GX8 | 0.18 | 3.27E -06 | 0.10 | 4.75 E-07 | nd | | ns | | RTE, RetroTransposable Element,Skipper GAG-PRO |
| DDB0238172 | DDB_G0267728 | Q55GC4 | 0.36 | 0.030 | 0.08 | 0.011 | nd | | nd | | uduA3, upreg. in dupA mutant close to UDPA1 |
| DDB0238173 | DDB_G0267726 | Q55GC5 | 0.36 | 0.017 | 0.13 | 0.009 | nd | | nd | | uduA1, upregulated in dupA mutant |
| DDB0191175 | DDB_G0285793 | P54657 | 0.23 | 0.001 | 0.27 | 0.002 | ns | | ns | | cadA, calcium-dependent cell adhesion molecule 1 |
| DDB0191525 | DDB_G0287587 | P54661 | 0.34 | 0.005 | 0.29 | 0.001 | 0.09 | 0.007 | 0.18 | 0.013 | smlA, small aggregate formation protein |
| DDB0230179 | DDB_G0273017 | Q558S7 | nd | | nd | | 0.20 | 0.003 | 0.14 | 0.001 | isocitrate lyase |
| DDB0219940 | DDB_G0275437 | P15649 | nd | | nd | | 0.09 | 0.079 | 0.08 | 0.001 | 7E, HssA/B-like prot. 27-late development |

**Table S2.** List of proteomics files available at the ProteomeXchange Consortium via the PRIDE partner repository.

**A. FOS-Veg_All replicates**

| | .RAW file name |
|---|---|
| control | SPY_5A_FOS_veg_1_25ul_1Mar24_1.raw |
| | SPY_5A_FOS_veg_1_25ul_1Mar24_2.raw |
| | SPY_5A_FOS_veg_1_25ul_1Mar24_3.raw |
| | SPY_KO_FOS_veg_2_5ul_22Apr24_1.raw |
| | SPY_KO_FOS_veg_2_5ul_22Apr24_2.raw |
| | SPY_KO_FOS_veg_2_5ul_22Apr24_3.raw |
| | SPY_KO_FOS_veg_2_25ul_27Apr24_1.raw |
| | SPY_KO_FOS_veg_2_25ul_27Apr24_2.raw |
| | SPY_KO_FOS_veg_2_25ul_27Apr24_3.raw |
| | SPY_KO_FOS_veg_rep2_1ul_21Jun24_1.raw |
| | SPY_KO_FOS_veg_rep2_1ul_21Jun24_2.raw |
| | SPY_KO_FOS_veg_rep2_1ul_21Jun24_3.raw |
| | SPY_KO_FOS_veg_rep3_0_75ul_24Jun24_1.raw |
| | SPY_KO_FOS_veg_rep3_0_75ul_24Jun24_2.raw |
| | SPY_KO_FOS_veg_rep3_0_75ul_24Jun24_3.raw |
| sample | AX3_4A_FOS_veg_1_5ul_1Mar24_1.raw |
| | AX3_4A_FOS_veg_1_5ul_1Mar24_2.raw |
| | AX3_4A_FOS_veg_1_5ul_1Mar24_3.raw |
| | AX3_FOS_veg_2_75ul_22Apr24_1.raw |
| | AX3_FOS_veg_2_75ul_22Apr24_2.raw |
| | AX3_FOS_veg_2_75ul_22Apr24_3.raw |
| | AX3_FOS_veg_1_5ul_27Apr24_1.raw |
| | AX3_FOS_veg_1_5ul_27Apr24_2.raw |
| | AX3_FOS_veg_1_5ul_27Apr24_3.raw |
| | AX3_FOS_veg_rep2_1ul_24Jun24_1.raw |
| | AX3_FOS_veg_rep2_1ul_24Jun24_2.raw |
| | AX3_FOS_veg_rep2_1ul_24Jun24_3.raw |
| | AX3_FOS_veg_rep3_0_55ul_24Jun24_1.raw |
| | AX3_FOS_veg_rep3_0_55ul_24Jun24_2.raw |
| | AX3_FOS_veg_rep3_0_55ul_24Jun24_3.raw |

PRIDE accession PXD056853
Results: AX3_SPY_FOS_veg_4_Bio_5_tech.mzid
Peaks: AX3_SPY_FOS_veg_4_Bio_5_tech.mzML

date of submission 15-Oct-24
Submission # 1-20241015-212944-1926934

**B. FOT-Veg_All replicate**s

| | .RAW file name |
|---|---|
| control | SPY_5B_FOT_veg_1_25ul_1Mar24_1.raw |
| | SPY_5B_FOT_veg_1_25ul_1Mar24_2.raw |
| | SPY_5B_FOT_veg_1_25ul_1Mar24_3.raw |
| | SPY_KO_FOT_veg_2ul_23Apr24_1.raw |
| | SPY_KO_FOT_veg_2ul_23Apr24_2.raw |
| | SPY_KO_FOT_veg_2ul_23Apr24_3.raw |
| | SPY_KO_FOT_veg_2_5ul_29Apr24_1.raw |
| | SPY_KO_FOT_veg_2_5ul_29Apr24_2.raw |
| | SPY_KO_FOT_veg_2_5ul_29Apr24_3.raw |
| | SPY_KO_FOT_veg_rep2_0_75ul_21Jun24_1.raw |
| | SPY_KO_FOT_veg_rep2_0_75ul_21Jun24_2.raw |
| | SPY_KO_FOT_veg_rep2_0_75ul_21Jun24_3.raw |
| | SPY_KO_FOT_veg_new_r3_1ul_03Jul24_1.raw |
| | SPY_KO_FOT_veg_new_r3_1ul_03Jul24_2.raw |
| | SPY_KO_FOT_veg_new_r3_1ul_03Jul24_3.raw |
| sample | AX3_4B_FOT_veg_1_5ul_1Mar24_1.raw |
| | AX3_4B_FOT_veg_1_5ul_1Mar24_2.raw |
| | AX3_4B_FOT_veg_1_5ul_1Mar24_3.raw |
| | AX3_FOT_veg_2ul_23Apr24_1.raw |
| | AX3_FOT_veg_2ul_23Apr24_2.raw |
| | AX3_FOT_veg_2ul_23Apr24_3.raw |
| | AX3_FOT_veg_2_5ul_29Apr24_1.raw |
| | AX3_FOT_veg_2_5ul_29Apr24_2.raw |
| | AX3_FOT_veg_2_5ul_29Apr24_3.raw |
| | AX3_FOT_veg_rep2_0_75ul_21Jun24_1.raw |
| | AX3_FOT_veg_rep2_0_75ul_21Jun24_2.raw |
| | AX3_FOT_veg_rep2_0_75ul_21Jun24_3.raw |
| | AX3_FOT_veg_new_r3_1ul_03Jul24_1.raw |
| | AX3_FOT_veg_new_r3_1ul_03Jul24_2.raw |
| | AX3_FOT_veg_new_r3_1ul_03Jul24_3.raw |

PRIDE accession PXD056857
Results:
AX3_SPY_FOT_veg_3_5_Bio_5_tech.mzid
Peaks:
AX3_SPY_FOT_veg_3_5_Bio_5_tech.mzML

date of submission 16-Oct-24
Submission # 1-20241016-142622-1926934

**C. FOS-Slug_All replicates**

| | .RAW file name |
|---|---|
| control | SPY_2A_FOS_slug_4ul_4Mar24_1.raw |
| | SPY_2A_FOS_slug_4ul_4Mar24_2.raw |
| | SPY_2A_FOS_slug_4ul_4Mar24_3.raw |
| | SPY_KO_FOS_slug_3_5ul_18Apr24_1.raw |
| | SPY_KO_FOS_slug_3_5ul_18Apr24_2.raw |
| | SPY_KO_FOS_slug_3_5ul_18Apr24_3.raw |
| | SPY_KO_FOS_slug_4_5ul_25Apr24_1.raw |
| | SPY_KO_FOS_slug_4_5ul_25Apr24_2.raw |
| | SPY_KO_FOS_slug_4_5ul_25Apr24_3.raw |
| | SPY_KO_FOS_slug_new_r1_1_75ul_17Jul24_1.raw |
| | SPY_KO_FOS_slug_new_r1_1_75ul_17Jul24_2.raw |
| | SPY_KO_FOS_slug_new_r1_1_75ul_17Jul24_3.raw |
| sample | AX3_1A_FOS_slug_2ul_29Feb24_0.raw |
| | AX3_1A_FOS_slug_6ul_5Mar24_1.raw |
| | AX3_1A_FOS_slug_6ul_5Mar24_2.raw |
| | AX3_1A_FOS_slug_6ul_5Mar24_3.raw |
| | AX3_FOS_slug_3_25ul_18Apr24_1.raw |
| | AX3_FOS_slug_3_25ul_18Apr24_2.raw |
| | AX3_FOS_slug_3_25ul_18Apr24_3.raw |
| | AX3_FOS_slug_3_5ul_25Apr24_1.raw |
| | AX3_FOS_slug_3_5ul_25Apr24_2.raw |
| | AX3_FOS_slug_3_5ul_25Apr24_3.raw |

AX3_FOS_slug_new_r1_1_75ul_18Jul24_1.raw
AX3_FOS_slug_new_r1_1_75ul_18Jul24_2.raw
AX3_FOS_slug_new_r1_1_75ul_18Jul24_3.raw

PRIDE accession PXD056903
Results: AX3_SPY_FOS_slug_4_reps_original.mzid
Peaks:
AX3_SPY_FOS_slug_4_reps_original.mzML

date of submission 16-Oct-24
Submission # 1-20241016-163729-1926934

**D. FOT-Slug_All replicates**

| | .RAW file name |
|---|---|
| control | SPY_2B_FOT_slug_4_5ul_4Mar24_1.raw |
| | SPY_2B_FOT_slug_4_5ul_4Mar24_2.raw |
| | SPY_2B_FOT_slug_4_5ul_4Mar24_3.raw |
| | SPY_KO_FOT_slug_3_25ul_19Apr24_1.raw |
| | SPY_KO_FOT_slug_3_25ul_19Apr24_2.raw |
| | SPY_KO_FOT_slug_3_25ul_19Apr24_3.raw |
| | SPY_KO_FOT_slug_4_5ul_26Apr24_1.raw |
| | SPY_KO_FOT_slug_4_5ul_26Apr24_2.raw |
| | SPY_KO_FOT_slug_4_5ul_26Apr24_3.raw |
| | SPY_KO_FOT_slug_new_r1_1_5ul_09Jul24_1.raw |
| | SPY_KO_FOT_slug_new_r1_1_5ul_09Jul24_2.raw |
| | SPY_KO_FOT_slug_new_r1_1_5ul_09Jul24_3.raw |
| sample | AX3_1B_FOT_slug_3_5ul_5Mar24_1.raw |
| | AX3_1B_FOT_slug_3_5ul_5Mar24_2.raw |
| | AX3_1B_FOT_slug_3_5ul_5Mar24_3.raw |
| | AX3_FOT_slug_4ul_19Apr24_1.raw |
| | AX3_FOT_slug_4ul_19Apr24_2.raw |
| | AX3_FOT_slug_4ul_19Apr24_3.raw |
| | AX3_FOT_slug_4_75ul_26Apr24_1.raw |
| | AX3_FOT_slug_4_75ul_26Apr24_2.raw |
| | AX3_FOT_slug_4_75ul_26Apr24_3.raw |
| | AX3_FOT_slug_new_r1_1_25ul_09Jul24_1.raw |
| | AX3_FOT_slug_new_r1_1_25ul_09Jul24_2.raw |
| | AX3_FOT_slug_new_r1_1_25ul_09Jul24_3.raw |

PRIDE accession PXD056866
Results:
AX3_SPY_FOT_slug_4_original_prep_runs.mzid
Peaks:
AX3_SPY_FOT_slug_4_original_prep_runs.mzML

date of submission 16-Oct-24
Submission # 1-20241016-182054-1926934

**E. FOS/T-Veg_Spy overexpression replicates**

| | .RAW file name | |
|---|---|---|
| SPY_KO (3) | SPY_5A_FOS_veg_1_25ul_1Mar24_1.raw | FOS |
| | SPY_5A_FOS_veg_1_25ul_1Mar24_2.raw | |
| | SPY_5A_FOS_veg_1_25ul_1Mar24_3.raw | |
| | SPY_5B_FOT_veg_1_25ul_1Mar24_1.raw | FOT |
| | SPY_5B_FOT_veg_1_25ul_1Mar24_1.raw | |
| | SPY_5B_FOT_veg_1_25ul_1Mar24_1.raw | |
| SPY_OE (2) | sample_6A_FOS_T_3_5ul_8Mar24_1.raw | FOS |
| | sample_6A_FOS_T_3_5ul_8Mar24_2.raw | |
| | sample_6A_FOS_T_3_5ul_8Mar24_3.raw | |
| | sample_6B_FOS_T_3_5ul_8Mar24_1.raw | FOT |
| | sample_6B_FOS_T_3_5ul_8Mar24_2.raw | |
| | sample_6B_FOS_T_3_5ul_8Mar24_3.raw | |
| AX3 (1) | AX3_4A_FOS_veg_1_5ul_1Mar24_1.raw | FOS |
| | AX3_4A_FOS_veg_1_5ul_1Mar24_2.raw | |
| | AX3_4A_FOS_veg_1_5ul_1Mar24_3.raw | |
| | AX3_4B_FOT_veg_1_5ul_1Mar24_1.raw | FOT |
| | AX3_4B_FOT_veg_1_5ul_1Mar24_1.raw | |
| | AX3_4B_FOT_veg_1_5ul_1Mar24_1.raw | |

PRIDE accession PXD057018
Results: AX3_SPY_OE_KO_veg_cells_FOS_T_rep1.mzid
Peaks: AX3_SPY_OE_KO_veg_cells_FOS_T_rep1.mzML

date of submission 21-Oct-24
Submission # 1-20241021-153215-1926934
